# Seven mutations unlock strict synthetic methylotrophy in engineered *Pseudomonas putida*

**DOI:** 10.64898/2026.07.20.739708

**Authors:** Òscar Puiggené, Martina Fricano, Riccardo Rossi, Luc F. M. Jansen, Emre Özdemir, Se Hyeuk Kim, Christina Lenhard, Elsayed T. Mohamed, Stefano Donati, Jochen Förster, Viji Kandasamy, Adam M. Feist, Enrico Orsi, Pablo I. Nikel

## Abstract

Methanol is a reduced, soluble one-carbon (C_1)_ feedstock for sustainable bioproduction, but converting this potential into robust microbial growth remains difficult. Several synthetic C_1_ assimilation routes depend on autocatalytic cycles, whose operation requires coordinated control of redox balance, toxic intermediates, substrate regeneration, and host regulation. Here, we implemented the serine-threonine cycle (STC) in the soil bacterium *Pseudomonas putida* and used growth-coupled selection with adaptive laboratory evolution (ALE) to transition from mixotrophic C_1_ incorporation to strict methylotrophy. The evolved strain grew with methanol as the sole carbon and energy source under atmospheric CO_2_ with a doubling time of ca. 40 h. Whole-genome sequencing, reverse genetics, biosensors, isotope labelling, and comparative RNA sequencing showed that evolution repeatedly targeted native pyrroloquinoline quinone (PQQ)-dependent methanol oxidation, membrane-bound transhydrogenase activity, glycine regeneration, STC enzyme balance, and global regulatory nodes. Additional ALE under glycine-methanol co-feeding increased growth rates and exposed further targets for improving cycle flux. These results establish *P. putida* as a chassis for strict synthetic methylotrophy and define actionable engineering routes toward C_1_ biomanufacturing.

**GRAPHICAL ABSTRACT:** 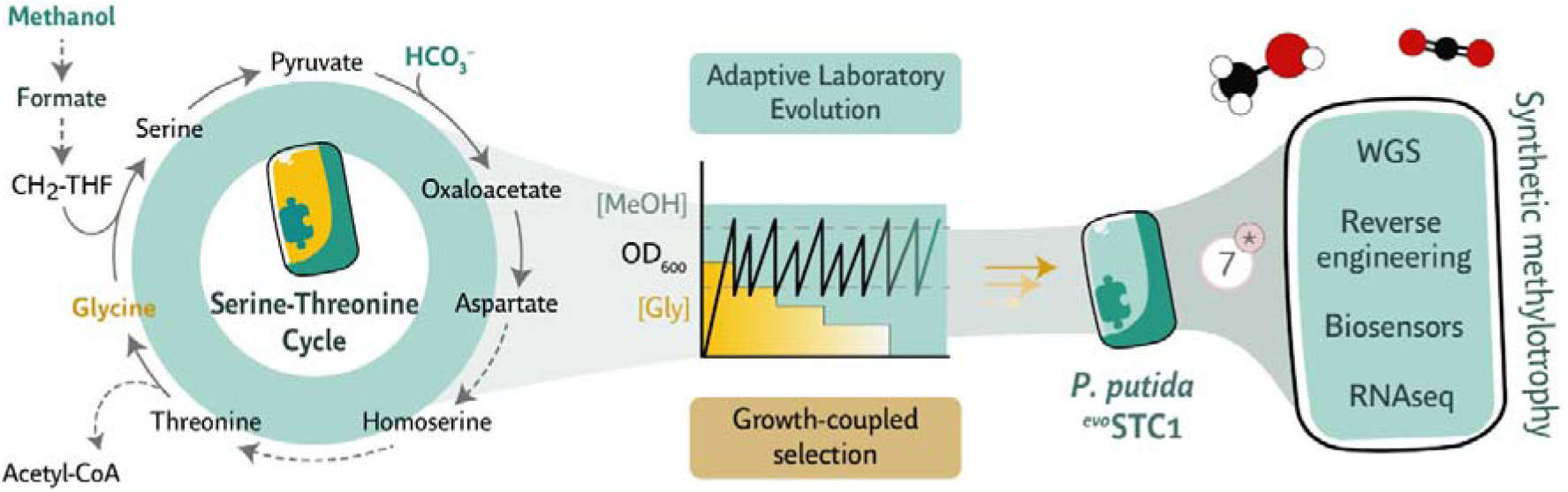

## 1. INTRODUCTION

Human-driven greenhouse gas emissions have accelerated climate change, threatening ecosystems and human well-being. Addressing this issue requires substantial emissions reductions, improved CO_2 c_apture technologies, development of a circular carbon (bio)economy, and a decisive shift toward renewable energy (Matthews et al., 2022; Rogelj et al., 2021). Within this broader transition, one-carbon (C_1)_ compounds, e.g., methanol and formate, have attracted increasing interest as alternative feedstocks for industrial bioproduction (Baumschabl et al., 2024; García and Galán, 2022; Gleizer et al., 2020; Tan et al., 2024). These substrates can be derived from CH_4_ or CO_2_ using renewable energy through (electro)chemical processes and do not compete with food and feed supply chains (Awogbemi and Desai, 2025; Liu et al., 2025; Lovat et al., 2025). Their soluble and reduced nature also circumvents gas-to-liquid transfer limitations, offering technical advantages over gas fermentation. Methanol is a particularly attractive feedstock because of its high reduction state, potential yields, and comparatively low toxicity (Park et al., 2024; Puiggené et al., 2025a).

The main limitation for C_1_ biomanufacturing is the conversion of this feedstock potential into robust microbial growth and production platforms. Chemical C_1_ conversion routes have advanced, yet they remain constrained by narrow product ranges, harsh reaction conditions, limited specificity, and high energy demands (Hepburn et al., 2019; Liu et al., 2016). Biological systems offer a complementary route, enabling production of value-added compounds under milder conditions (He et al., 2026; Hu et al., 2024; Lee et al., 2025). However, conventional industrial microbes, e.g., *Escherichia coli* and Saccharomyces cerevisiae, lack natural C_1_ assimilation pathways (Sanford and Woolston, 2022), whereas native methylotrophs remain difficult to engineer for broad product synthesis at high yield, despite substantial progress towards their domestication (Claassens et al., 2016; Liu et al., 2020; Ørsted et al., 2026; Whitaker et al., 2015). Efforts in synthetic methylotrophy therefore aim to combine the metabolic logic of C_1_ assimilation with the genetic tractability and industrial familiarity of established microbial chassis.

Recent metabolic engineering efforts have enabled synthetic methylotrophy in model organisms, taking advantage of well-characterized metabolism and efficient genetic tools (Kim et al., 2020; Orsi et al., 2023; Reiter et al., 2024; Zhong et al., 2023). Nevertheless, the field still faces a substantial implementation gap. Many synthetic C_1_ pathways are thermodynamically plausible and can function under mixotrophic or auxotrophic selection regimes, yet few support robust, strict C_1_-dependent growth. Autocatalytic pathway architectures are particularly difficult to transplant because growth requires coordinated balancing of redox metabolism, cofactor supply, toxic intermediate handling, substrate regeneration, and native regulatory programs. Rational design provides the initial pathway architecture, whereas adaptive laboratory evolution (ALE) can tune fluxes, relieve regulatory conflicts, and reveal otherwise hidden physiological constraints (Fernández-Cabezón et al., 2019). Interpreting such evolutionary outcomes requires pathway-specific selection schemes, isotope tracing, genome sequencing, biosensors, and systems-level analyses. Closing the implementation gap in synthetic C_1_ metabolism therefore requires iterative strategies that combine rational pathway construction, evolution-based optimization, and mechanistic dissection of evolved genotypes and phenotypes (Puiggené et al., 2025a; Reiter et al., 2024).

Expanding the range of chassis organisms is equally important. Among potential hosts, the Gram-negative soil bacterium *Pseudomonas putida* displays several traits that are advantageous for synthetic C_1_ assimilation, including high physicochemical stress tolerance (Calero and Nikel, 2019; de Lorenzo et al., 2024; Weimer et al., 2020) and native C_1_-related enzymatic activities (Turlin et al., 2023) that may help overcome metabolic limitations, e.g., substrate processing. Besides, *P. putida* benefits from an expanding genetic and genome-engineering toolset (Martínez-García and de Lorenzo, 2024; Volke et al., 2023) and displays strong potential as a host for production of fine chemicals, including new-to-nature products (Calero et al., 2020; Nikel and de Lorenzo, 2018; Wirth and Nikel, 2021). Although *P. putida* does not naturally assimilate C_1_ compounds, this bacterium encodes several dehydrogenases capable of generating reducing equivalents from methanol, formaldehyde, and formate, including the pyrroloquinoline quinone (PQQ)-dependent alcohol dehydrogenases PedE and PedH, which can act as methanol dehydrogenases (MeDHs) (Turlin et al., 2023). The bacterium also possesses native pyruvate and phosphoenolpyruvate carboxylases, which can support efficient carboxylation toward oxaloacetate (Chavarría et al., 2012; Nikel et al., 2015; 2021). Previous studies from our laboratory established *P. putida* as a chassis for strict formate and methanol assimilation through the reductive glycine pathway (Turlin et al., 2022; 2025). More recently, synthetic serine cycles promoted methanol incorporation under mixotrophic conditions (Puiggené et al., 2025b). The adaptable metabolism and environmental robustness of *P. putida* therefore make this host well suited for advancing synthetic C_1_ assimilation from partial pathway activity to strict methylotrophy.

The serine-threonine cycle (STC) holds high potential for implementing synthetic methylotrophy (Puiggené et al., 2025b). This O_2-_insensitive pathway integrates methanol or formate assimilation with CO_2_ fixation and produces acetyl-coenzyme A (acetyl-CoA) under ambient CO_2_ levels without carbon loss (Bar-Even, 2016; Wenk et al., 2024). The STC also displays favorable thermodynamics and harnesses endogenous reactions widely present across microbial hosts (He et al., 2020; Puiggené et al., 2025a). Furthermore, the STC is among the few synthetic C_1_ assimilation cycles that have been fully implemented in a heterotrophic host. The seminal work of Wenk et al. (2024) demonstrated formatotrophic growth in engineered *E. coli*, although the resulting strain had limited performance for industrial-scale applications. These observations indicate that the STC is a suitable biochemical architecture for C_1_-trophy, but successful implementation still depends on extensive tuning of host physiology. More broadly, synthetic C_1_ metabolism has relied heavily on ALE, yet evolved strains often accumulate multiple mutations whose individual and combined effects are difficult to resolve, leaving a persistent challenge in linking genotype, pathway function, and C_1_-trophic phenotype.

Here, we engineered *P. putida* for strict methanol-based growth using growth-coupled selection and ALE. The output of this campaign is an engineered *P. putida* strain capable of strict growth on methanol and atmospheric CO_2_ with the STC as the C_1_ assimilation architecture. Seven mutations were sufficient to mediate synthetic methylotrophy, and their effects were dissected through reverse genetics, biosensors, isotope labelling patterning, whole-genome sequencing, and systems-level transcriptomics. In parallel, ALE under low-glycine and high-methanol conditions identified metabolic and regulatory bottlenecks that limit growth performance. The analysis shows that methanol oxidation, transhydrogenase activity, glycine regeneration, and regulatory rewiring must be coordinated to establish synthetic methylotrophy. Together, the experiments reported here define the evolutionary and physiological steps required to convert STC activities in engineered *P. putida* from mixotrophic C_1_ incorporation into strict methylotrophic growth.

## 2. RESULTS

### 2.1. *13C-methanol tracing experiments reveal limited carbon propagation through the STC in engineered* P. putida

Our previous work identified the STC as the synthetic serine-cycle topology with the highest potential for enabling methylotrophy in *P. putida* (Puiggené et al., 2025b), and we demonstrated STC activity through relief of L-serine auxotrophy in a mixotrophic setup. Here, we sought to move beyond simple auxotrophic complementation and establish full methylotrophic growth through ALE. STC-harboring strains supported C_1_-dependent growth when glucose, acetate, pyruvate, or glycerol were supplied as co-substrates, but failed to grow when glycine was supplied as the main carbon source together with formate or methanol (data not shown). To examine the metabolic constraints underlying this phenotype, we analyzed ^13^C-methanol incorporation in strain STC1. This strain contains a genomically integrated C_1_-assimilation module and constitutively overexpresses ltaE, encoding L-threonine aldolase for glycine regeneration, and *yiaY/PP_2683*, encoding regulators of native PQQ-dependent MeDHs (Puiggené et al., 2025b). Strain STC1 is also auxotrophic for serine and 5,10-methylene-tetrahydrofolate because of the Δ*serA* ΔΔ*gcv* deletions (Bushin et al., 2026) and carries additional modifications to prevent carbon efflux from the cycle and to suppress biofilm formation (Fig. S1).

We next examined whether enhanced serine aldolase activity could increase methanol-derived carbon incorporation. For this purpose, we compared the parental STC configuration with an enhanced serine-threonine cycle (eSTC) variant expressing a mutated E. *coli L*-threonine aldolase, *^Ec^ltaE*. This variant carries the C188Y substitution, hereafter termed *^Ec^ltaE*\*, previously shown to improve the aldolase activity and increase growth parameters of selection strains co-fed with methanol (Puiggené et al., 2025b; Schann et al., 2024). *^Ec^ltaE*\* was expressed from the low-copy-number vector pSEVA621 and compared with an empty-vector control in P. putida STC1. Cultures were grown in minimal salt medium (MSM) with 125 mM ^13^C-labelled methanol, 20 mM glucose, and 10 nM lanthanum chloride (LaCl_3)_ to promote activity of the native PedH MeDH. The expected (theoretical) and observed (experimental) labelling patterns are shown in Fig. 1a and 1b, respectively. Glycine remained unlabeled, whereas serine displayed complete single-carbon incorporation. The eSTC variant showed only a modest increase in carbon labelling relative to the parental STC strain, likely because glucose reduced the demand for C_1_ assimilation under these conditions.

**Figure 1.**
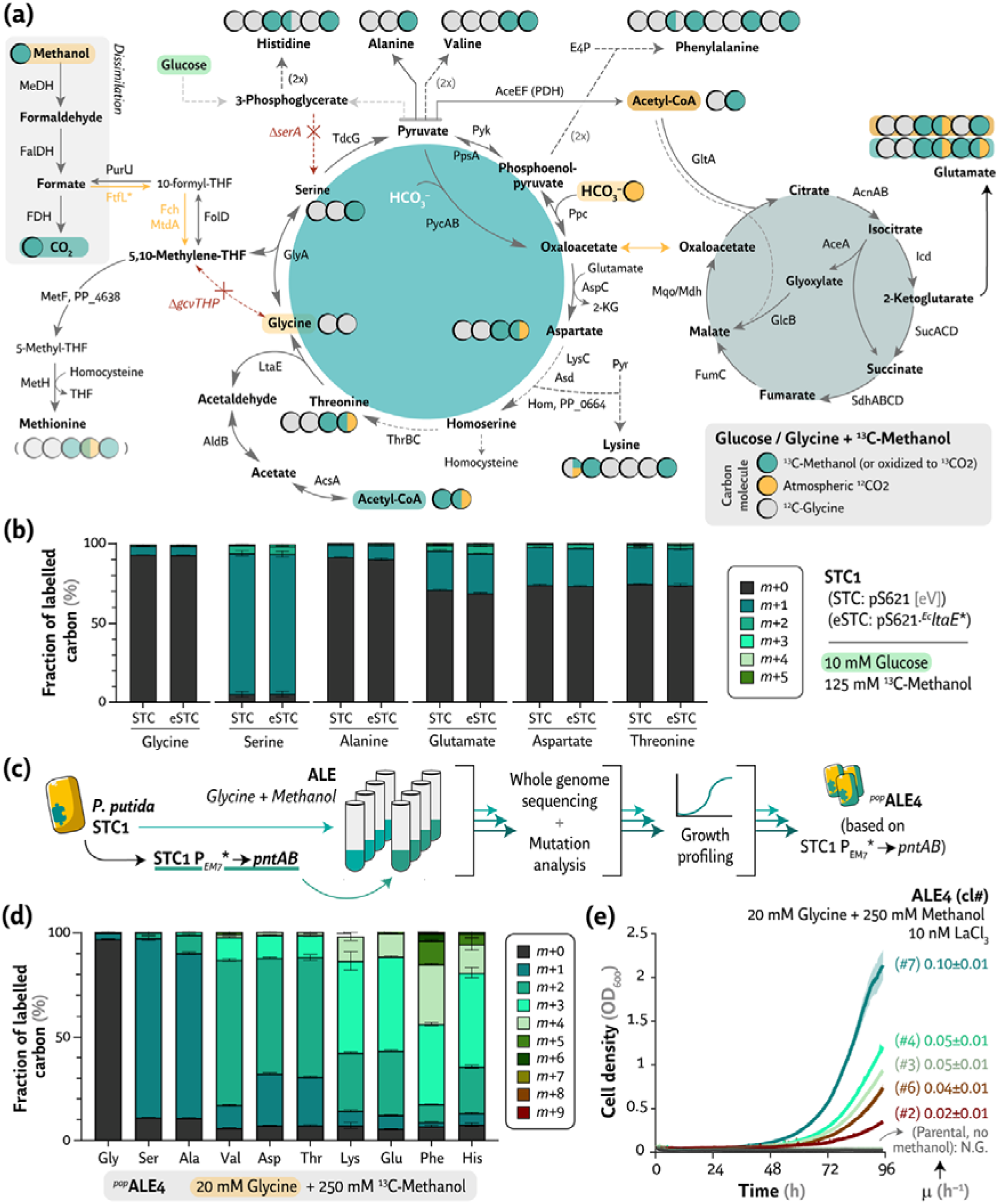
Isotope labeling proves methanol assimilation and identifies putative metabolic bottlenecks limiting synthetic methylotrophy. **(a)** Metabolic map showing the ^13^C-labeling profile following growth on ^13^C-methanol with either glycine or glucose. Atmospheric ^12^CO_2_ fixation is indicated in yellow, whereas oxidation of labeled methanol and subsequent reassimilation reactions via formate are shown in teal. Unlabeled carbon atoms are displayed in gray. **(b)** Amino acid labeling patterns associated with the synthetic serine cycle (STC) in the STC1 strain carrying either the biosensor plasmid pS621·^Ec^ltaE* or the empty vector control (eSTC vs. STC). Cultures were grown in MSM supplemented with 10 mM glucose, 125 mM ^13^C-methanol, and 10 nM LaCl_3._ (**c**) PALE adaptive laboratory evolution (ALE) strategy used to enable growth of STC1 and STC1 P_EM7*_→pntAB on glycine and methanol. The mutational profiles of the evolved populations are provided in Table S2. (**d**) Amino acid labeling patterns associated with the synthetic serine cycle in the evolved population *^pop^*ALE4 grown in MSM supplemented with 20 mM glycine, 250 mM ^13^C-methanol, and 10 nM LaCl_3._ (**e**) Growth profiles, indicated as optical density at 600 nm (OD_600)_, of individual clones isolated from *^pop^*ALE4 grown in MSM supplemented with 20 mM glycine, 250 mM methanol, and 10 nM LaCl_3._ The parental strain (unevolved P. putida STC1 P_EM7*_→pntAB) and no-methanol controls are included. Specific growth rates (μ, h^−1^) are indicated on the right side of the panel. In all cases, the results represent averages ± standard deviation from at least three independent biological experiments. Key abbreviations: Acetyl-CoA, acetyl-coenzyme A; Gly, glycine; Ser, serine; Ala, alanine; Val, valine; Asp, aspartate; Thr, threonine; Lys, lysine; Glu, glutamate; Phe, phenylalanine; His, histidine; eV, empty vector; and N.G., no growth. Further information on enzyme abbreviations, reactions, and functions in panel (**a**) can be found in Table S1.

To identify metabolic constraints associated with STC operation, we then examined labelling in amino acids closely connected to cycle intermediates, namely alanine, glutamate, aspartate, and threonine (Fig. 1b). Alanine showed limited labelling, with approximately 10% of the *m*+1 fraction, whereas glutamate, aspartate, and threonine showed similar labelling profiles, close to *m*+1 = 25%. These patterns are consistent with two possible scenarios: carboxylation of glucose-derived pyruvate using fully oxidized C_1_ units, or carboxylation of a labelled C_3 c_ompound using atmospheric CO_2_ (Fig. 1a). The low level of labelling in alanine, which derives directly from pyruvate, supports the first scenario as the most likely.

Overall, ^13^C-methanol tracing substantiated C_1_ incorporation into specific STC-linked intermediates but showed limited propagation of labelled carbon beyond growth-coupled metabolites. This profile indicates substantial methanol oxidation together with restricted autocatalysis (i.e., cyclic operation) or insufficient glycine regeneration. These observations indicated that STC activity was detectable but poorly propagated through central metabolism, prompting us to use ALE to increase flux through the synthetic assimilation cycle.

### 2.2. ALE drives methanol-derived carbon flux during glycine-supported STC growth

We implemented a two-stage evolutionary strategy: first, adaptation to growth in MSM supplemented with glycine and methanol, followed by progressive selection for full methylotrophy. The glycine-methanol stage was designed to favor biomass formation through the initial assimilatory reactions of the STC, corresponding to Module 1. Progression to strict methylotrophic growth would then require glycine regeneration and sustained autocatalysis (Fig. 1a). Both stages followed a PALE (*Pathway Activation of Latent Enzymes*) ALE framework (Guzmán et al., 2019), in which the concentration of the auxiliary carbon source was gradually reduced to force increased reliance on the engineered pathway.

In addition to strain STC1, we included a derivative constitutively expressing *pntAB* from the evolved *P_EM7_*\* promoter, hereafter referred to as STC1 *P_EM7_*\*→*pntAB*. This design was intended to alleviate the NADPH limitation previously observed during synthetic C_1 a_ssimilation schemes. Deletion of aceA, which had been used to increase NAD(P)H and ATP availability during implementation of the reductive glycine pathway, was unsuitable here because the glyoxylate shunt is required for STC autocatalysis. Through this shunt, the two acetyl-CoA molecules generated by the STC can replenish cycle intermediates, several of which also serve as amino acid precursors for biomass formation (Fig. 1a).

Acetate was initially selected as an additional supporting substrate together with glycine and methanol. Parsimonious flux balance analysis indicated that acetate metabolism more closely resembles the flux distribution predicted for STC-based growth than glucose metabolism (Fig. S2). In *P. putida*, acetate strongly activates the glyoxylate shunt, a prerequisite for STC autocatalysis. The in silico simulations also predicted lower pyruvate dehydrogenase activity under acetate-grown conditions than under glucose-grown conditions, thereby reducing carbon leakage from the cycle.

After serial passages with decreasing acetate concentrations, evolved populations grew in MSM supplemented with 20 mM glycine, 250 mM methanol, and 10 nM LaCl_3._ These populations were subjected to whole-genome sequencing and growth profiling (Fig. 1c). Key high-confidence mutations are summarized in Table S2 and discussed below. Among the evolved lineages, *^pop^*ALE4, derived from STC1 P*_EM7_*\*→*pntAB*, was selected for further analysis because it displayed improved growth while lacking an obvious causative mutation that could explain growth with glycine as the main carbon source. We then performed isotope labelling and isolated 48 individual clones for phenotypic characterization (Fig. 1d and 1e). The best-performing isolate, *^pop^*ALE4 clone 7, reached μ ∼ 0.1 h^−1^ and final optical density at 600 nm (OD_600)_ = 2 (Fig. 1e). Other isolates from the same population showed distinct growth behaviors, suggesting that *^pop^*ALE4 may contain subpopulations with complementary adaptations.

To determine whether *^pop^*ALE4 growth depended on methanol assimilation, we repeated ^13^C-labelling experiments with this evolved population (Fig. 1d), focusing on amino acids derived from STC-linked intermediates. Serine displayed single-carbon labelling, whereas glycine remained unlabeled, as expected. Alanine, derived from pyruvate transamination, showed predominantly single-carbon incorporation. Strikingly, 55% of aspartate appeared as the *m*+2 isotopomer, indicating that methanol-derived carbon was oxidized to CO_2_ (or HCO_3–_) and subsequently reassimilated through carboxylation reactions into oxaloacetate, the direct precursor of aspartate. Additional amino acids, including valine, threonine, lysine, glutamate, phenylalanine, and histidine, showed higher-than-expected labelling, even after accounting for reassimilation of completely oxidized methanol. Taken together, the labelling patterns indicate that methanol-derived carbon contributed at least one-third of biomass carbon, based on serine and aspartate labelling, and that STC autocatalysis was active but still suboptimal. Since STC operation requires assimilation of one C_1_ unit together with one glycine molecule, growth on glycine imposes a stronger demand for C_1_ assimilation than growth with glucose as co-substrate. Higher autocatalytic flux should therefore increase labelling across replenished cycle intermediates. To dissect the regulatory and metabolic basis of *^pop^*ALE4 adaptation, we next performed transcriptomic analysis.

### 2.3. Transcriptomics captures regulatory rewiring in ^pop^ALE4

To dissect the adaptation trajectory toward growth on glycine and methanol, we profiled the transcriptome of *^pop^*ALE4 and its parental strain STC1 P*_EM7*_→pntAB* under regimes combining glycine and/or acetate, methanol, and rare earth elements (REEs). The resulting dataset offered some mechanistic resolution of the key STC-related adaptations (Fig. S3). Expression of the ped cluster and other C_1_-dissimilatory genes was slightly lower in *^pop^*ALE4 than in the parental strain, suggesting attenuated methanol oxidation capacity. By contrast, expression of the membrane-bound transhydrogenase *pntAB* genes was modestly elevated, consistent with increased demand for NADPH balancing during C_1_ assimilation.

Transcriptional patterns in central carbon metabolism genes also showed signs of rewiring. Tricarboxylic acid cycle genes were generally downregulated in *^pop^*ALE4, whereas glyoxylate shunt components remained expressed at robust levels, matching the expected metabolic requirements of STC operation. Notably, *tdcG-III* was strongly induced in response to glycine supplementation, while *glyA* homologues showed no comparable transcriptional increase (Fig. S3). This pattern may reflect an uneven distribution of flux through the cycle or a bottleneck in glycine-to-serine conversion. Overall, transcriptomics revealed regulatory adjustments consistent with STC activity but did not identify a single transcriptional mechanism explaining growth of *^pop^*ALE4 on glycine and methanol. Because transcriptomics did not resolve a single dominant mechanism behind glycine-supported growth, we next increased the selection pressure and evolved STC-harboring cells toward strict methanol dependence.

### 2.4. ALE enables strict STC-dependent methylotrophy at low growth rates

Following adaptation to glycine and methanol in several evolved populations, we initiated the second stage of the PALE-ALE strategy (Fig. 2a). Five STC1 populations pre-adapted to these substrates were propagated using automated serial passaging. The goal was to progressively reduce the concentration of the auxiliary substrate once cultures exceeded the growth threshold of their parental passages, indicating improved fitness (Fig. 2a). In this regime, glycine concentrations were decreased when growth improved but were increased again when population performance declined (Fig. S4). Populations after each passage were also evaluated for their ability to sustain strict methylotrophic growth.

**Figure 2.**
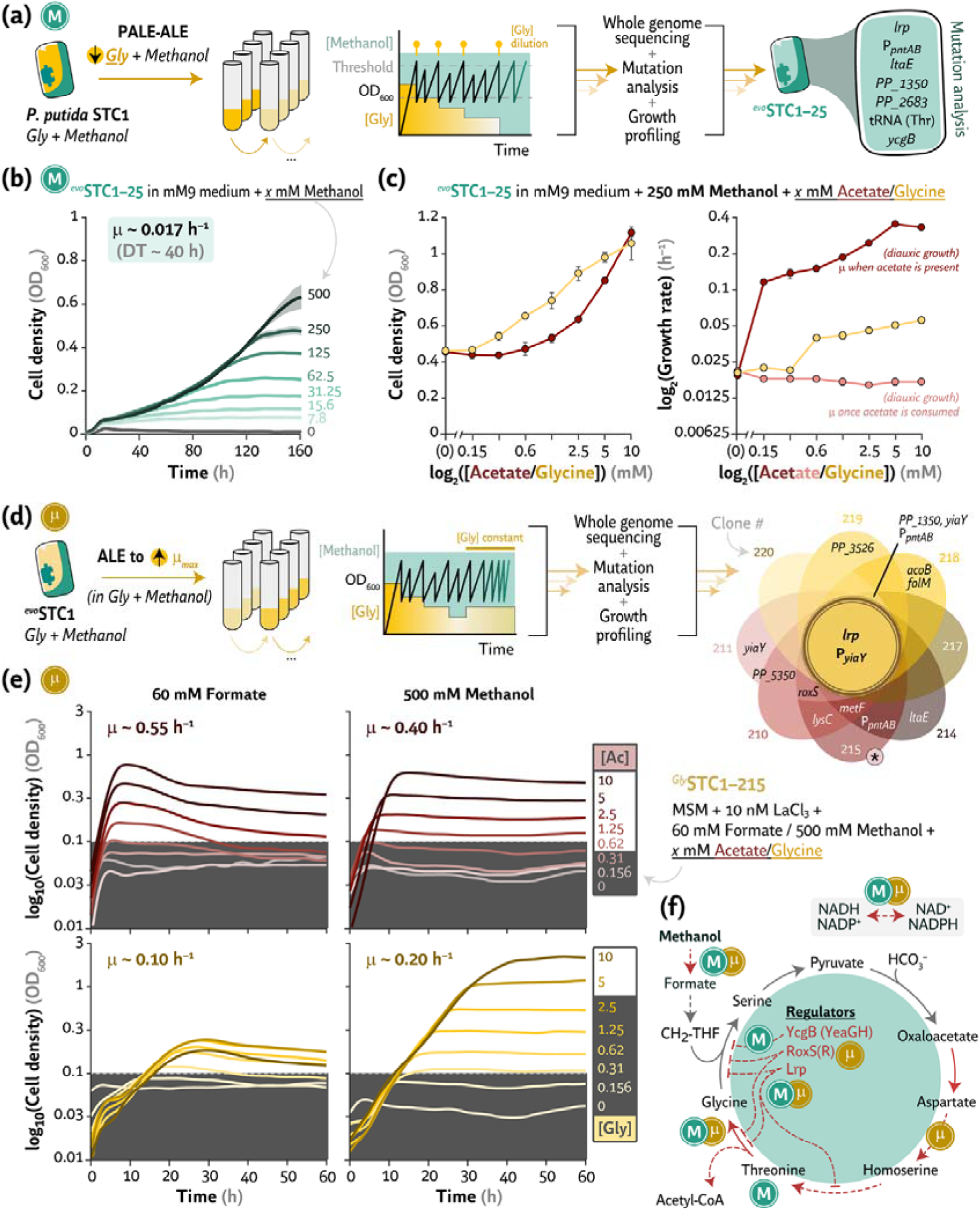
ALE promotes methylotrophic growth and enhances methanol assimilation by reducing the requirement for glycine as a secondary carbon source. (**a**) ALE strategy used to establish full methylotrophy from the STC1 strain by progressively decreasing the glycine concentration as methanol assimilation improved. Following clone isolation, whole genome sequencing, and mutational analysis, genes carrying consensus mutations in the resulting strain ^evo^STC1-25 are indicated. (**b**) Growth profiles of P. putida ^evo^STC1-25 in mM9 medium (see also Fig. S6a) supplemented with increasing methanol concentrations. (**c**) Final cell densities and specific growth rates of ^evo^STC1-25 grown in mM9 medium supplemented with 250 mM methanol and increasing concentrations of acetate or glycine as secondary carbon sources. Acetate resulted in diauxic growth (see also Fig. S6c); therefore, specific growth rates for both growth phases are shown in red and pink, corresponding to growth before and after acetate depletion, respectively. (**d**) ALE strategy used to increase the growth rate of the STC1 strain previously evolved for growth on glycine and methanol in panel (**a**) while maintaining a constant glycine concentration to further improve methanol assimilation. Following clone isolation, whole genome sequencing, and mutational analysis, genes carrying consensus mutations shared among the isolated clones are indicated. Based on growth performance, clone 215, designated ^Gly^STC1-215, was selected for further characterization. (**e**) Growth profiles of P. putida ^Gly^STC1-215 in mM9 medium supplemented with either 60 mM formate or 500 mM methanol and increasing concentrations of acetate or glycine. Specific growth rates during exponential growth are indicated. (**f**) Metabolic map highlighting recurrent mutations identified in both ALE experiments, including mutations affecting the transhydrogenase encoded by *pntAB* and the regulatory genes ycgB, roxS, and lrp. Key abbreviations: DT, doubling time; Gly, glycine; Ac, acetate; and CH_2-_THF, 5,10-methylenetetrahydrofolate.

Since specific growth rates remained low throughout this PALE-ALE campaign, we reasoned that other nutritional limitations could be at play. Therefore, we shifted to a M9 minimal medium supplemented with Wolfe’s vitamin mix, SL7 trace elements, and a REE mixture, hereafter referred to as modified M9 medium (mM9). Wolfe’s vitamin mix contains riboflavin, nicotinic acid, calcium D-(+)-pantothenate, vitamin B_12_, biotin, folic acid, and other cofactors. In this new formulation, the REE includes not only lanthanum but also heavier lanthanides, which may stimulate native MeDH activities (Wehrmann et al., 2017). These supplements were included to better support cell viability and growth under the stress imposed by prolonged selection for methylotrophy.

After extended cultivation, one population from the PALE-ALE experiment sustained growth on 500 mM methanol as the sole carbon source (Fig. 2a-c and Fig. S4). Several clones from this population were isolated and sent for whole-genome sequencing. Virtually all clones shared the same genotype, hereafter named ^evo^STC1-25, defined by seven mutations affecting *lrp*, the *pntAB* promoter, *ltaE, PP_1350, PP_2683*, the L-threonine tRNA *PP_t73*, and *ycgB* (Fig. 2a, Table 1, and Table S3). An extended record of this evolution experiment is also available in the ALEdb database (Phaneuf et al., 2019). The first five mutations occurred in all sequenced clones, whereas the *PP_t73* and *ycgB* mutations were absent from one clone. The absence of the *ycgB* mutation was most likely offset by the concomitant appearance of the *yeaG*^Y463*^ mutation (Table S3), although whether other mutations compensated for the missing L-threonine tRNA substitution remains unclear.

**Table 1.** Mutations identified in the fully methylotrophic evolved strain ^evo^STC1-25, including the affected genes, functional consequences, and putative adaptive significance.

| Gene | Mutation reported<sup>a</sup><br>(clone count) | Gene function | Mutational significance |
| --- | --- | --- | --- |
| *lrp* | D12Y (26/26) | Leucine-responsive protein | Deregulation of the serine-glycine-threonine node<sup>b</sup> |
| *P<sub>pntAB</sub>* | –117C→T (26/26) | (Promoter of) Membrane-bound transhydrogenase complex genes | Increased production of transhydrogenase PntAB |
| *ltaE* | A73T (26/26) | Low specificity L-threonine aldolase | Balancing fluxes towards glycine regeneration |
| *PP_1350* | T261P (26/26) | Histidine kinase | Cofactor biosynthesis deregulation<sup>b</sup> |
| *PP_2683* | 17+A (26/26) | Hybrid histidine kinase | Reducing formaldehyde toxicity |
| *PP_t73* | 12A→G (25/26) | tRNA for L-threonine (ACA codon) | Decreasing threonine depletion and node balancing<sup>b</sup> |
| *ycgB* | 394(CGGAACAGGTA)1→2 (25/26) | Serine/threonine kinase | Modulating post-translational regulation of STC enzymes<sup>b</sup> |

In the following sections, we examine the putative effects of the mutations present in ^evo^STC1-25. First, we characterized growth under strict methylotrophic conditions and in cultures supplemented with acetate or glycine as secondary carbon sources (Fig. 2b and 2c).

### 2.5. Growth profiling of P. putida ^evo^STC1-25 across partial and strict C_1_-trophic regimes

Clone ^evo^STC1-25 grew under strict methylotrophic conditions, with methanol serving as the only carbon and energy source. We cultivated *P. putida* ^evo^STC1-25 in 96-well plates containing mM9 medium with increasing methanol concentrations, ranging from 8 mM to 500 mM (Fig. 2b). At 250 mM methanol, cultures reached OD_600 ∼_ 0.5, whereas supplementation with 500 mM methanol supported OD_600 >_ 0.6. Final cell densities increased with methanol concentration, while cultures without methanol showed no detectable growth. This result substantiated the methylotrophic nature of this isolate, indicating that the remaining medium components did not support appreciable biomass formation (Fig. 2b). ^13^C-isotope labelling of ^evo^STC1-25 grown in the presence of ^13^C-methanol supported this interpretation, with 85-90% labelled carbon in the measured amino acids. The remaining fraction is consistent with incorporation of atmospheric ^12^CO_2_ through endogenous carboxylases, including phosphoenolpyruvate and pyruvate carboxylases (Fig. S5). Growth remained slow, with μ ∼ 0.017 h^−1^, corresponding to a doubling time = 40 h.

Because this ALE campaign used a medium composition that differs from standard P. putida cultivation conditions, we compared full mM9 medium with the same formulation lacking either SL7 trace elements or Wolfe’s vitamins, and MSM supplemented with either REEs or LaCl_3 (_Fig. S6). Across methanol concentrations, full mM9 medium supported the highest final cell density at 500 mM methanol, consistently reaching OD_600 >_ 0.6. Removing trace amounts of Wolfe’s vitamins or SL7 caused only minor differences, and specific growth rates remained essentially unchanged across the mM9 medium variants, with μ ∼ 0.017 h^−1^ (Fig. S6a). MSM supported lower final cell densities and slightly lower growth rates than mM9 medium. LaCl_3 p_erformed similarly to, or slightly better than, the REE mixture (Fig. S6a). We therefore retained mM9 medium for subsequent fully C_1_-trophic experiments.

Having established the evolved methylotrophic phenotype, we next assessed whether *P. putida* ^evo^STC1-25 could grow on formate in mM9 medium, since formate lies downstream of methanol oxidation. Interestingly, increasing formate concentrations supported weaker growth than methanol, with OD_600 <_ 0.2 and μ ∼ 0.008 h^−1^ (Fig. S6b). This reduced performance in formate cultures likely reflects the lower energy and reducing-equivalent yield of this more oxidized C_1_ substrate. Higher formate concentrations may also impose toxicity and further restrict growth.

Higher cell densities and growth rates are often essential for bioprocess viability. Since either methanol or formate promoted low cell densities when supplemented as the only carbon substrate, we evaluated *P. putida* ^evo^STC1-25 under mixotrophic conditions using acetate or glycine as secondary carbon sources (Fig. 2c). Acetate concentrations above 0.5 mM and glycine concentrations above 0.15 mM substantially increased final cell densities. Acetate triggered diauxic growth, with a pronounced increase in the initial specific growth rate followed by a return to basal methylotrophic rates once acetate was depleted (Fig. 2c and Fig. S6c). Glycine supplementation produced a near-linear increase in OD_600 a_nd a weaker but consistent increase in μ values (Fig. 2c). This response suggests that glycine regeneration remains a limiting factor in ^evo^STC1-25. The linear trend is also consistent with the inherent STC stoichiometry, as glycine assimilation is coupled to methanol incorporation in an approximately 1:1 ratio. These observations identified glycine regeneration as a persistent limitation in ^evo^STC1-25 and motivated a second ALE campaign designed to improve growth with glycine-methanol co-feeding.

### 2.6. ALE under glycine-methanol co-feeding selects faster growth and higher STC flux

Because glycine regeneration appeared to limit strict methylotrophic growth at higher rates, we used ALE again to target this bottleneck. Building on the previous PALE-ALE experiments performed at low glycine concentrations in the presence of methanol, we ran serial passaging at constant glycine concentrations of ca. 2.65-3 mM (Fig. 2d and Fig. S4). Specific growth rates increased over time under these conditions (Fig. S7), after which clones were isolated from all populations and subjected to whole-genome sequencing and growth profiling (Fig. S8). These isolates carried diverse mutational profiles, summarized in Fig. 2d and Table S4. Among them, clone ^Gly^STC1-215 displayed the highest growth rate in mM9 medium supplemented with 500 mM methanol and 2.65 mM glycine (Fig. S8), and was therefore selected for detailed characterization.

We quantified growth of ^Gly^STC1-215 across increasing glycine or acetate concentrations as secondary carbon sources, using mM9 medium supplemented with either 60 mM sodium formate or 500 mM methanol and 10 nM LaCl_3 (_Fig. 2e). Under these conditions, *P. putida* ^Gly^STC1-215 reached higher final OD_600 v_alues and specific growth rates than ^evo^STC1-25, particularly in glycine-formate cultures. Still, glycine and formate supported relatively low final cell densities, likely because formate supplies less reducing power and energy than methanol. This limitation was not observed when acetate was provided, consistent with its capacity to generate energy through the tricarboxylic acid cycle. Acetate-methanol cultures reached specific growth rates approximately twice as fast as those observed with glycine, although final OD_600 v_alues were lower (Fig. 2e).

We also performed isotope labelling analysis of strain ^Gly^STC1-215 grown in mM9 medium supplemented with 10 mM unlabeled glycine, 500 mM ^13^C-methanol, and 10 nM LaCl_3._ These data were compared with labelling profiles from strain STC1, grown with glucose as co-substrate because STC1 does not grow on glycine, and from *^pop^*ALE4 (Fig. 1d and Fig. S5a). Strain ^Gly^STC1-215 showed higher degrees of amino acid labelling than either STC1 or *^pop^*ALE4, with up to 30% of STC-associated amino acids being fully labelled. Fully-labelled STC-derived amino acids were negligible in STC1 and *^pop^*ALE4; a pattern that indicates that ALE under glycine-methanol co-feeding increased flux toward glycine regeneration.

Across the ALE campaigns, recurrent mutations targeted several key functional layers of the engineered methylotrophic phenotype (Fig. 2f). These included genes involved in native methanol oxidation systems, the membrane-bound transhydrogenase, STC enzymes, and regulatory proteins. We therefore examined how recurrent mutations in these functional layers shaped STC flux, starting with LtaE, the enzyme directly linking L-threonine metabolism to glycine regeneration.

### 2.7. LtaE mutations tune glycine-regeneration flux during STC evolution

LtaE is a central enzyme of the STC architecture. In previous work, overexpression of ltaE increased flux from the tricarboxylic acid cycle, through oxaloacetate, toward glycine regeneration (Puiggené et al., 2025b). This design used the constitutive P*_trc_* _p_romoter and the canonical SEVA ribosome-binding site (RBS). Because rational metabolic engineering rarely identifies optimal expression or activity levels a *priori*, ALE can help tune pathway flux during selection. In our methylotrophic ALE campaigns, the engineered ltaE promoter and RBS were not affected. Instead, the ltaE open reading frame (ORF) was targeted twice: an A73T change arose during selection for strict methylotrophy, whereas the Y134N mutation appeared during selection for faster growth under glycine-methanol conditions. The A73T mutation maps close to the Mg^2+^ and Cl^−^ ions at the LtaE monomer interface, while the Y134N modification may influence pyridoxal phosphate cofactor binding in the enzyme (Fig. 3a).

**Figure 3.**
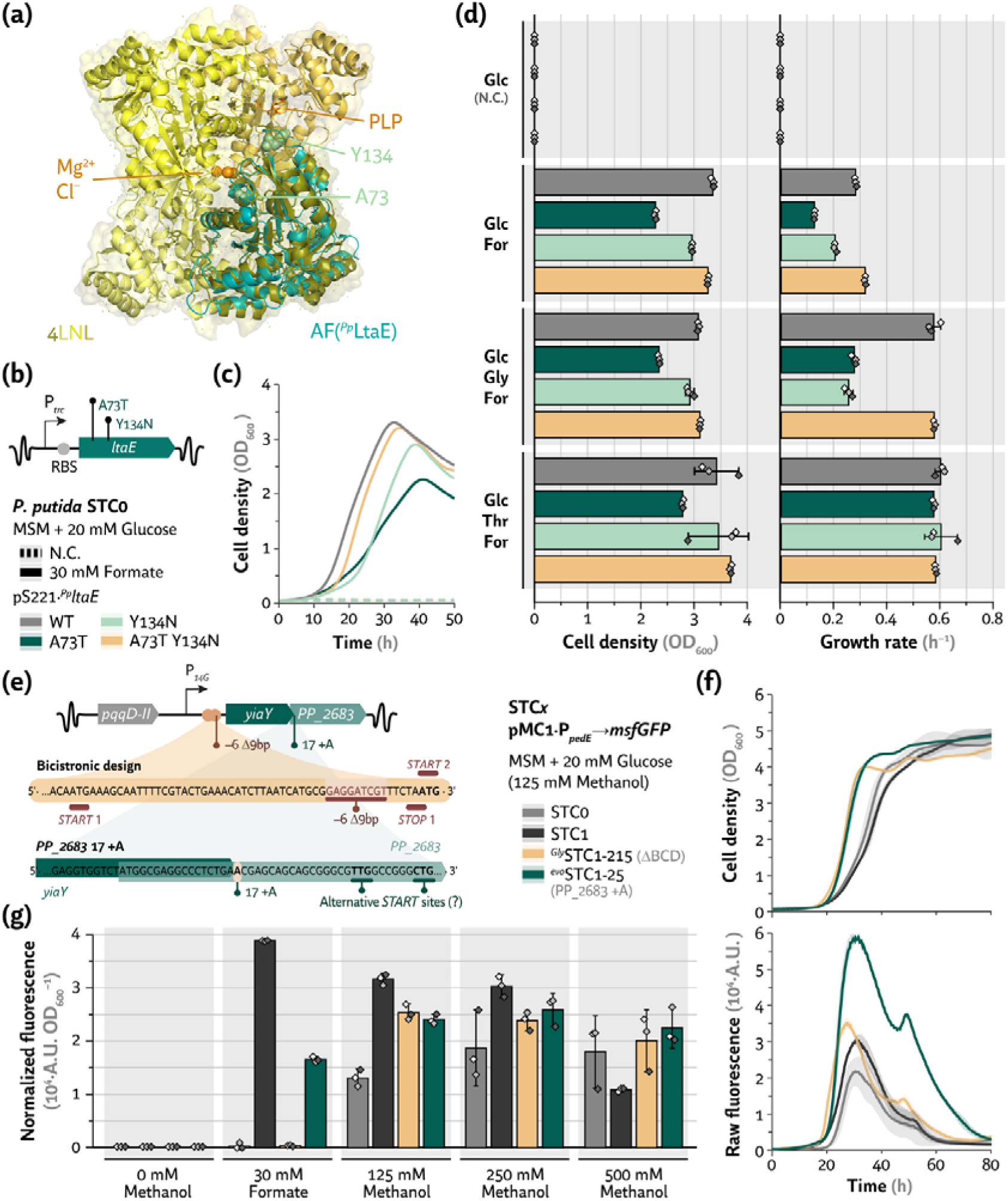
ALE balances key metabolic nodes to improve methylotrophic growth. Characterization of mutations affecting the L-threonine aldolase encoded by ltaE (**a-d**) and the ped cluster regulator genes yiaY and PP_2683, which control PQQ-dependent methanol dehydrogenase activity (**e-g**). (**a**) AlphaFold (AF)-predicted structure of P. putida LtaE superimposed on the crystal structure of Escherichia coli LtaE (PDB: 4LNL). Magnesium (Mg^2+^) and chloride (Cl^−^) ions, the pyridoxal phosphate cofactor, and the mutated residues are highlighted. (**b**) Schematic representation of the A73T and Y134N mutations identified by ALE in the ltaE coding sequence, which is constitutively expressed from the chromosome under the P_trc p_romoter and the SEVA ribosome binding site (RBS). (**c**) Growth profiles of the STC0 strain carrying vector pSEVA221 expressing the wild-type (WT), two single-mutant, or double-mutant P. putida ltaE variants. Cultures were grown in MSM supplemented with 20 mM glucose in the presence, or absence (N.C., negative control) of 30 mM formate. (**d**) Maximum cell density (OD_600)_ and specific growth rate of the strains shown in panel (**c**) grown in MSM supplemented with 20 mM glucose (Glc) and, where indicated, 30 mM formate (For), 10 mM glycine (Gly), or 10 mM L-threonine (Thr). (**e**) Schematic representation of the genomic region containing the ped cluster regulator genes yiaY and PP_2683, highlighting the recurrent mutations identified during ALE. A partial deletion of BCD10 occurred in ^Gly^STC1-215, whereas a frameshift mutation in PP_2683 was identified in all ^evo^STC1-25 clones. (**f**) Growth and raw fluorescence profiles of strains STC0, STC1, ^Gly^STC1-215, and ^evo^STC1-25 carrying the biosensor plasmid pMC1·P_pedE→_msfGFP. Cultures were grown in MSM supplemented with 20 mM glucose and 125 mM methanol in the absence of rare earth elements to promote pedE expression. (**g**) Maximum normalized fluorescence (A.U. OD_600_^−1^) measured in MSM supplemented with 20 mM glucose and the indicated concentrations of methanol or formate. Data represents mean values ± standard deviation from three biological replicates; except for error bars in panels (**d**) and (**g**), which indicate 95% confidence intervals. Pairwise comparisons with non-overlapping error bars correspond to P < 0.05.

To characterize the physiological consequences of these substitutions, we constructed pSEVA221·*^Pp^ltaE* plasmids expressing the wild-type (WT) *ltaE* allele or the *ltaE*^A73T^ and ltaE^Y134N^ variants in the naïve strain STC0. In this context, we adopted strain STC0 to avoid confounding effects from overexpression of the *ped* cluster regulators encoded by *yiaY* and *PP_2683*, which were observed to be recurrently targeted during our ALE campaigns (Fig. 2). A double-mutant variant carrying both substitutions was also included to assess potential epistasis (Fig. 3b-d).

Growth profiling of strain STC0 expressing wild-type ltaE, the single mutants, or the double mutant was performed in MSM supplemented with 20 mM glucose and 30 mM formate. Both single mutations reduced flux through the L-threonine aldolase reaction, as indicated by only partial relief of the serine auxotrophy and lower final cell densities and growth rates relative to the wild-type allele (Fig. 3c and 3d). Supplementation with glycine or L-threonine did not fully restore growth, particularly for the strain carrying the A73T variant selected during strict methylotrophy. The double mutant did not show a synergistic defect, and the corresponding engineered strain had a growth pattern generally resembling the wild-type profile (Fig. 3c and 3d).

These results suggest that ALE selected LtaE variants that rebalance flux through the threonine biosynthetic branch rather than simply maximizing glycine regeneration. Excessive LtaE activity may drain L-threonine needed for protein synthesis, creating a trade-off between STC turnover and amino acid availability. Because threonine biosynthesis is also tightly controlled by allosteric regulation, increased L-threonine availability may further restrict upstream pathway flux, consistent with the lower OD_600 o_bserved the engineered strain bearing the LtaE^A73T^ variant. Having examined mutations affecting glycine regeneration, we next turned to the methanol oxidation node, where ALE repeatedly targeted the engineered regulators controlling the native ped cluster.

### 2.8. Mutations in yiaY and PP_2683 tune ped cluster expression during methylotrophic ALE

To promote PQQ-MeDH activity, we previously identified that upregulation of the *ped* cluster regulators *yiaY* and *PP_2683* increases expression of the genes encoding the endogenous PQQ-dependent methanol dehydrogenases, *pedE* or *pedH*, depending on REE availability (Puiggené et al., 2025b; Puiggené and Nikel, 2026). Although this strategy improves growth parameters upon methanol exposure, recurrent mutations in this locus indicate that strong regulator expression can become disadvantageous, particularly under relaxed (low selection) conditions. Consistent with this view, each sequenced clone carried a distinct mutation in the engineered promoter, RBS, or ORF of yiaY or PP_2683 (Fig. 2a and 2d). To assess the consequences of these changes, we examined the capacity of strains ^evo^STC1-25 and ^Gly^STC1-215 to activate pedE expression in the absence of lanthanides. These clones carried different alterations in this regulatory region: *P. putida* ^evo^STC1-25 contained a frameshift (+A) near the *N*-terminus of *PP_2683* at position 17, whereas *P. putida* ^Gly^STC1-215 had a 9-bp deletion in the bicistronic design BCD10 used as the RBS to drive overexpression of ped genes in strain STC1 (Fig. 3e).

We compared these evolved clones with parental strains STC0 and STC1, with strain STC0 carrying the intact ped cluster and strain STC1 serving as the ancestor of the evolved lineages. The four strains were transformed with the biosensor plasmid pMC1·P*_pedE_*→*msfGFP*, in which the native *pedE* promoter drives expression of the monomeric superfolder GFP (*msfGFP*) gene (Fig. 3e-g). Cultures were grown in MSM with 20 mM glucose and increasing methanol concentrations, from 125 to 500 mM, or with 30 mM formate as a negative control. The resulting growth profiles, raw msfGFP fluorescence, and OD_600-_normalized fluorescence are shown in Fig. 3f and 3g.

Relative to their parental counterparts, both *P. putida* ^evo^STC1-25 and ^Gly^STC1-215 displayed lower OD_600-_normalized msfGFP fluorescence at 125-250 mM methanol (Fig. 3g). At 500 mM methanol, growth defects likely caused by increased toxicity altered this pattern, and strain STC1 no longer produced the highest normalized reporter fluorescence. The evolved clones also maintained ped cluster expression until stationary phase, whereas expression in parental strains STC0 and STC1 declined shortly after mid-exponential phase (Fig. 3f). The expression profile of *P. putida* ^evo^STC1-25 remained closer to that of strain STC1, suggesting that the frameshift in *PP_2683* did not fully abolish regulator functionality. One plausible explanation is the potential translation from downstream *start* sites, which would reduce translational efficiency and tune ped cluster activation in response to methanol. By contrast, the BCD10 deletion detected in strain ^Gly^STC1-215 shifted expression closer to that observed in strain STC0, particularly without methanol or with formate, where fluorescence remained consistently low (Fig. 3g).

Overall, mutations in the ped cluster regulators appear to rebalance the methanol oxidation node by reducing peak ped cluster expression while extending activity through exponential phase. This lower expression may reduce toxicity caused by excessive formaldehyde formation, which endogenous C_1_-dissimilation routes may not detoxify efficiently under methylotrophic selection.

### 2.9. Increased transhydrogenase activity is key for C_1 a_ssimilation in synthetic hosts

Transhydrogenases help maintain intracellular redox balance by catalyzing reversible transfer of reducing equivalents between the NADH/NAD^+^ and NADPH/NADP^+^ pools (Partipilo et al., 2026). This function is particularly important during C_1_-assimilation pathway implementation, including the reductive glycine pathway and serine cycles, which often impose high demand for NADPH-dependent reductive reactions (Bar-Even, 2016). Several studies establishing synthetic C_1_-trophy in *E. coli* and *P. putida* have consistently recovered ALE mutations that further increase expression of genes encoding transhydrogenase systems, especially the membrane-bound, proton-gradient-dependent *pntAB* complex (Kim et al., 2020; Kim et al., 2023; Satanowski et al., 2020; Turlin et al., 2025; Wenk et al., 2024).

Although we previously attempted direct *pntAB* overexpression through the P_*EM7*\*_→*pntAB* construct, the ALE campaign toward strict methylotrophy was designed to let strain STC1 adapt from its native regulatory capacity. Mutations in the *pntAB* region appeared in all clones evolved for strict methylotrophy, including ^evo^STC1-25, and in most clones selected for higher growth rates under glycine-methanol selection, including the ^Gly^STC1-215 lineage (Fig. 2a and 2d, and Tables S3 and S4). Notably, ^Gly^STC1 clones lacking pntAB-region mutations, such as ^Gly^STC1-210 and ^Gly^STC1-211, carried mutations in PP_5350, encoding an RpiR-like transcriptional regulator (Table S4). Among the recurrent *pntAB* promoter mutations, a C→T substitution at position –117 appeared consistently across ALE campaigns (Fig. 4a).

**Figure 4.**
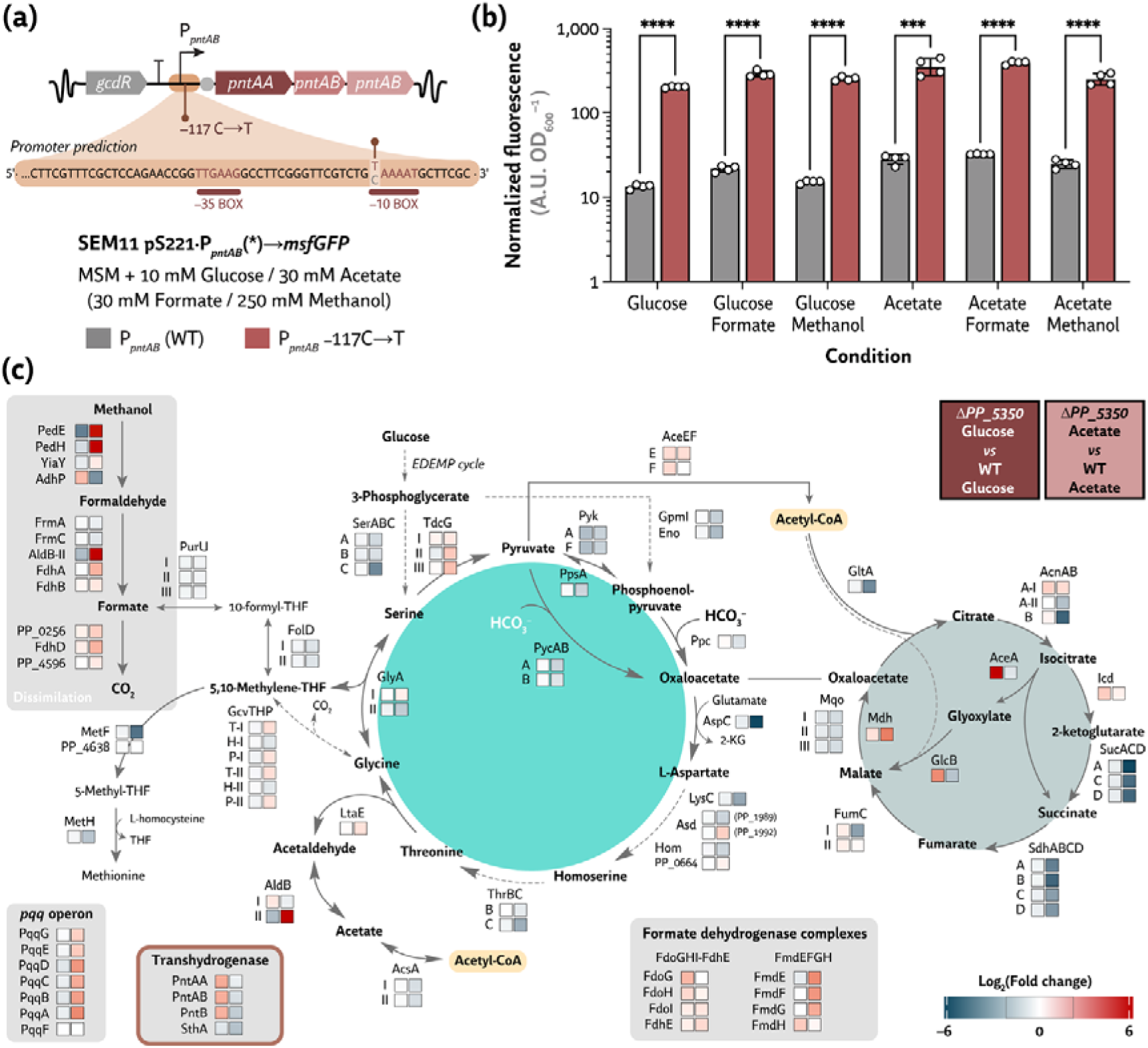
Transhydrogenases play a key role in methanol assimilation in engineered *P. putida*. (**a**) During ALE, a –117C→T mutation arose in the endogenous *pntAB* promoter, probably affecting the –10 promoter element. (**b**) Normalized fluorescence (A.U. OD_600–_^1^) of SEM11 cells carrying the biosensor plasmid pSEVA221·P_pntAB→_msfGFP containing either the evolved or the wild-type promoter. Cultures were grown in MSM supplemented with 10 mM glucose or 30 mM acetate and, where indicated, 30 mM formate or 250 mM methanol. Data represents mean values ± standard deviation from four biological replicates. Statistical significance is indicated as *** (P < 0.001) and **** (P < 0.0001) calculated using multiple unpaired t-tests, corrected by the Holm-Šídák method. (**c**) RNA-seq analysis of strain KT2440 ΔPP_5350 grown on acetate or glucose relative to the wild-type (WT) strain. Genes directly or indirectly associated with the synthetic serine cycle are highlighted. Three biological replicates were analyzed for each strain and growth condition and log_2 o_f the fold change is given. Further information on enzyme abbreviations, reactions, and functions in panel (**c**) can be found in Table S1.

To determine the effect of this mutation, we constructed a set of biosensor plasmids carrying either the wild-type or evolved P*_pntAB_* promoter upstream of an msfGFP reporter in the pSEVA621 backbone. These constructs were transformed into parental strain SEM11, and the resulting strains were grown in MSM with either 10 mM glucose or 30 mM acetate, with 30 mM formate or 250 mM methanol added where indicated (Fig. 4b). OD_600-_normalized msfGFP fluorescence was consistent across conditions and showed that the evolved promoter increased reporter expression by ca. one order of magnitude, corresponding to 10-to 20-fold higher activity than the wild-type P*_pntAB_* promoter (Fig. 4b). This result substantiates a high level of *pntAB* as a requisite towards implementation of C_1_-trophy, consistent with previous results across synthetic C_1_ assimilation pathways. We therefore asked whether evolved lineages lacking direct *pntAB* promoter mutations increased transhydrogenase gene expression through alternative regulatory changes.

### 2.10. Mutations in PP_5350 suggest an indirect route to *pntAB* upregulation

PP_5350 is an RpiR-family transcriptional regulator linked to repression of the glyoxylate shunt genes (Lim et al., 2021; Lim et al., 2022). We previously observed frameshift mutations in *PP_5350* during adaptation to glycine-methanol growth (Table S2) and during ALE for increased growth rates under glycine-methanol selection (Fig. 2d and Table S4). Since a previous study showed that a 9-base-pair deletion in PP_5350 enhances glyoxylate shunt activity (Lim et al., 2021), we performed RNA-seq analysis comparing wild-type *P. putida* KT2440 and the deletion mutant *ΔPP_5350* (Volke et al., 2021a). This comparison was used to assess both the expected effect on the glyoxylate shunt and broader consequences for central carbon metabolism (Fig. 4c).

Wild-type *P. putida* and the Δ*PP_5350* mutant were grown in MSM supplemented with either glucose or acetate, and samples were harvested during exponential phase. As expected, glyoxylate shunt genes were strongly upregulated in Δ*PP_5350* during growth on glucose (Fig. 4c). Loss of PP_5350 also increased expression of the ped cluster and the associated PQQ biosynthesis operon under acetate conditions. By contrast, genes encoding L-aspartate aminotransferase and tricarboxylic acid cycle components, including 2-oxoglutarate dehydrogenase and succinate dehydrogenase, were strongly downregulated relative to the wild-type control during acetate growth. Interestingly, the membrane-bound transhydrogenase operon *pntAB* was also upregulated under glucose conditions, whereas several formate dehydrogenase genes showed increased expression during growth on acetate (Fig. 4c). Apart from the expected derepression of the glyoxylate shunt, glucose-grown cells showed only limited transcriptional changes.

These data are consistent with a possible role for PP_5350 loss in reinforcing glyoxylate shunt activity, which could support the autocatalytic flux required for STC operation. However, mutations in PP_5350 consistently appeared only in evolved clones that lacked substitutions in the *pntAB* promoter (Fig. 2d and Table S4). This pattern suggests that PP_5350 inactivation may instead serve as an indirect route to increase expression of the membrane-bound transhydrogenase. Because PP_5350 loss likely causes broader regulatory and metabolic effects, direct mutations in the *pntAB* promoter appear to be the more efficient evolutionary solution for increasing *pntAB* expression. Beyond redox balancing and glyoxylate-shunt regulation, ALE also targeted broader transcriptional control, prompting us to examine the contribution of Lrp to STC performance.

### 2.11. Lrp inactivation has limited phenotypic impact on STC operation under mixotrophic conditions

The leucine-responsive regulatory protein Lrp is a highly conserved global regulator in bacteria. Lrp assembles as a tetramer of dimers, with each monomer containing an α/β C-terminal domain and an N-terminal helix-turn-helix DNA-binding domain (Fig. 5a). This regulator senses changes in nutrient availability and coordinates programs linked to virulence, motility, and nutrient acquisition (Ziegler and Freddolino, 2021). Although its full regulatory scope remains unresolved, Lrp is recognized as a master regulator of the feast-or-famine response in *E. coli* and may play a similar role in pseudomonads (González et al., 2018; Schmidt et al., 2022; Zhi et al., 1999). In *E. coli*, Lrp has been proposed to influence up to one-third of all transcriptional regulatory pathways (Kroner et al., 2019). Alignment of *P. putida* Lrp with the extensively characterized E. coli homolog revealed substantial sequence conservation, with 60% identity and 78% positives, as well as strong structural similarity at the dimeric level. Superposition of the AlphaFold-predicted *P. putida* Lrp dimer with the quaternary structure of E. coli Lrp, PDB 2GQQ (de los Rios and Perona, 2007), yielded a root mean square deviation = 0.957 Å (Fig. 5a).

**Figure 5.**
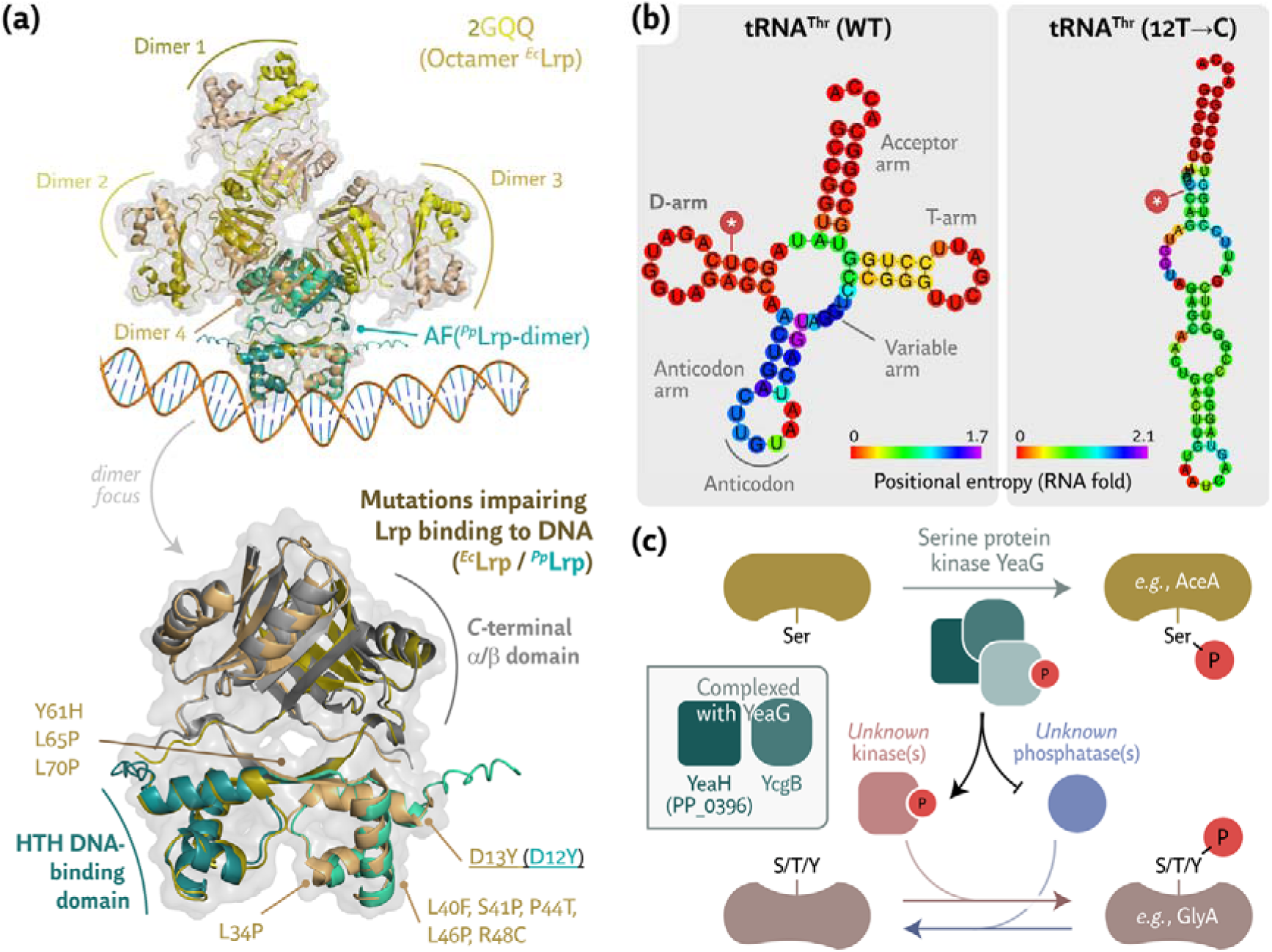
Structural modeling of mutations affecting Lrp, the L-threonine (Thr) tRNA, and the YcgB-YeaGH regulatory network. (**a**) Structural basis of Lrp octamer assembly and DNA binding. Top: Overview of the octameric E. coli Lrp complex (PDB: 2GQQ; yellow/tan) overlaid with an AlphaFold-predicted *P. putida* Lrp dimer (teal) bound to DNA. Bottom: Enlarged view of the dimer interface and the helix-turn-helix (HTH) DNA-binding domain. Key residues predicted to impair DNA binding when mutated (e.g., Y61H, L65P, L70P, and D13Y) are highlighted. Residue numbering corresponds to the equivalent positions in E. coli Lrp (tan) and *P. putida* Lrp (teal). (**b**) Secondary structure and positional entropy of the tRNA for L-threonine encoded by PP_t73. Predicted secondary structures of the wild-type and 12T→C mutant tRNA were generated using RNAfold from the ViennaRNA Package; colors indicate positional entropy, with separate legends shown for the wild-type and mutant structures. (**c**) Proposed YeaG-mediated phosphorylation pathway. The model depicts the protein kinase YeaG, together with YeaH (PP_0396) and YcgB, regulating downstream targets including AceA and GlyA through serine, threonine, and tyrosine phosphorylation, based on previous studies in E. coli. The complete regulatory network is expected to involve additional kinases and phosphatases that remain to be identified.

Several mutations are known to impair Lrp DNA-binding activity, as highlighted in Fig. 5a. Among them, the E. coli Lrp D13Y substitution is particularly relevant because it is homologous to the D12Y mutation detected in *P. putida* ^evo^STC1-25 (Tables 1 and S3). Although this residue lies in an helix-turn-helix region that does not directly contact DNA, substitutions in this segment can alter helix-turn-helix conformation and DNA-binding function (Platko and Calvo, 1993). These observations indicate that the D12Y mutation likely causes loss of Lrp function. In *E. coli*, lrp deletion increases expression of genes encoding key enzymes in the serine-threonine-glycine node, including *glyA, ltaE*, and *thrBC* (Cho et al., 2008; González et al., 2018). The emergence of D12Y in ^evo^STC1-25 may therefore reflect selection for increased flux toward L-threonine biosynthesis and glycine regeneration.

To assess the effect of lrp in *P. putida*, we deleted the gene from strains SEM11, the wild-type parent of STC1, STC0, and STC2, a pyruvate-decarboxylase-deficient STC1 derivative. The Δlrp mutants and their parental strains were grown in MSM supplemented with either 10 mM glucose or 30 mM acetate, together with 60 mM formate or 500 mM methanol. Methanol cultures were also supplemented with 10 nM LaCl_3._ Contrary to expectations, the *lrp* deletion did not confer a clear advantage or defect under mixotrophic conditions in any tested background, apart from minor, non-significant changes in lag phase (Fig. S9). Given the broad regulatory scope of Lrp, targeted overexpression of selected pathway components may offer a more controlled strategy to increase STC flux. On this background, we reasoned that full Lrp inactivation may benefit a fully methylotrophic background, but its pleiotropic effects could also constrain further gains in growth performance. Together, these targeted analyses show that strict methylotrophy emerged through coordinated changes in methanol oxidation, redox balancing, glycine regeneration, and global regulation, rather than through a single dominant mutation.

## 3. DISCUSSION

This work summarizes our efforts to establish a novel variant of the O_2-_insensitive, C_2-_yielding serine-threonine cycle (STC) as a thermodynamically favorable and genetically tractable design for C_1_ assimilation. By combining modular reconstruction, isotope tracing, ALE, and systems-level characterization, we established growth under fully methylotrophic conditions and identified metabolic and regulatory constraints repeatedly targeted during evolution campaigns. These constraints included intermediate leakage, insufficient glycine regeneration, altered enzyme expression, redox imbalance, and broader regulatory rearrangements. The recurrent nature of these targets across independent evolutionary stages emphasizes that the limiting steps were interconnected physiological pressures imposed by forcing an autocatalytic cycle into a non-native host rather than isolated defects. Together, these results expose the difficulty of engineering C_1_-assimilation cycles and the need to combine rational design, evolution, and mechanistic analysis to convert pathway activity into a growth-supporting phenotype.

*P. putida* is a robust and metabolically versatile chassis for synthetic C_1_ assimilation, partly because it encodes several alcohol dehydrogenases that can act on methanol. To increase native PQQ-MeDH activity, we previously overexpressed the *ped* cluster regulator genes *yiaY* and *PP_2683* constitutively (Puiggené et al., 2025b; Puiggené and Nikel, 2026). This strategy improved mixotrophic growth with glucose and enabled methanol assimilation at concentrations as low as 7.8 mM, indicating active PQQ-MeDH catalysis (Puiggené et al., 2025b). Because of their favorable thermodynamic properties, PQQ-MeDHs can oxidize methanol more efficiently at low concentrations than NAD(P)^+^-dependent MeDHs. However, the ped regulatory locus was repeatedly targeted during ALE. This pattern likely reflects the need to balance methanol oxidation with formaldehyde detoxification, since insufficient oxidation restricts carbon flux whereas excessive oxidation can increase formaldehyde toxicity through DNA and protein crosslinking (Roca et al., 2008; 2009). In the ^evo^STC1-25 and ^Gly^STC1-215 lineages, mutations in this region did not abolish methanol oxidation. Instead, they reduced peak ped cluster expression and extended activity across the exponential phase, a profile absent from parental strains. This sustained expression likely reflects adaptation to methanol as a carbon source rather than only as an electron donor, also indicating that optimal methanol oxidation depends on timing and dynamic control rather than maximal dehydrogenase induction.

ALE also tuned other metabolic branch points, including LtaE activity and PQQ-MeDH expression, while revealing functions that were present at insufficient basal levels. A clear example was the membrane-bound transhydrogenase PntAB. C_1_-assimilation pathways often impose high NADPH demands (Partipilo et al., 2026); in the STC, NADPH is required for glycine regeneration through L-threonine biosynthesis and for formate assimilation. Consistently, nearly all C_1_-trophy engineering efforts in heterotrophic bacterial hosts have recovered ALE mutations that increase transhydrogenase expression, particularly the H^+^-gradient-dependent *pntAB* complex (Kim et al., 2020; 2023; Satanowski et al., 2020; Turlin et al., 2025; Wenk et al., 2024). Here, a single mutation in the −10 region of the *pntAB* promoter increased expression by more than one order of magnitude. A loss-of-function mutation in *PP_5350* bypassed this promoter change but caused only a more modest transcriptional upregulation. These results indicate that native regulatory routes can raise *pntAB* expression, although the optimal expression level for a methylotrophic *P. putida* chassis remains to be resolved. This scenario also suggests that transhydrogenase activity should be treated as a core design variable in future C_1_-assimilation programs rather than as an accessory redox adjustment.

Several additional mutations in *P. putida* ^evo^STC1-25 affected poorly characterized regulatory or translational functions. The PP_t73 mutation affected one of the three *P. putida* tRNA L-threonine genes and altered a conserved nucleotide in the D-arm, a region required for proper tRNA folding and stability (Fig. 5b). PP_t73 decodes the ACA threonine codon, whereas PP_t03 and PP_t12 decode ACC and ACG, respectively. The mutation may reduce aminoacylation efficiency or render PP_t73 partly non-functional. Continued growth suggests that residual PP_t73 activity or wobble decoding by other tRNA isoacceptors sustains translation, although impaired ACA decoding may contribute to the long doubling time of strain ^evo^STC1-25. The low frequency (0.36%) of ACA codons in *P. putida*, compared with ACC at 3.1%, ACU at 0.47%, and ACG at 0.9% [www.kazusa.or.jp; (Arella et al., 2021)], may limit this burden. Direct biochemical analysis will be needed to define the effect of this mutation.

Other recurrently targeted loci included *yeaG/ycgB, roxS*, and *PP_1350*. YeaG is a PrkA-like serine/threonine kinase with an AAA^+^ module, and its gene colocalizes with yeaH and ycgB, which encode uncharacterized proteins. This conserved module has been linked to stress responses, nitrogen starvation, and nutrient utilization across several bacterial families (Figueira et al., 2015; Price et al., 2018). Perturbation of *yeaG* has also been associated with altered post-translational modification patterns in enzymes involved in C_1_ and central metabolism, including GlyA, AceA/B, SerA, Mdh, and PykA (Sultan et al., 2021). These links are relevant because GlyA, PykA, and the glyoxylate shunt shape STC flux, and GlyA activity can be directly modulated by post-translational modifications during carbon-source shifts (Brunk et al., 2018). In this context, *yeaG* or *ycgB* mutations could either directly increase STC flux, for example through GlyA, or reflect adaptation to glycine as a primary carbon source.

*PP_1350* encodes a sensor histidine kinase with no identified cognate response regulator in *Pseudomonas*. The PA4398 ortholog in *P. aeruginosa* PA14 affects swarming, biofilm formation, cyclic di-GMP metabolism, and siderophore-related genes (Strehmel et al., 2015). Three explanations may account for selecting mutations in PP_1350 during ALE. First, PP_1350 may help balance iron or REE uptake, since both metal classes can enter cells through overlapping transport systems and influence metalloenzyme activity. Second, its colocalization with *pduO* and *panE*, involved in vitamin B_12 a_nd pantothenate/CoA biosynthesis, raises the possibility that mutations reduce costly cofactor biosynthesis when these compounds are supplied in mM9 medium. Third, overexpression of yiaY could promote non-native interactions among histidine kinases, making PP_1350 mutations part of a broader rewiring of two-component signaling. These hypotheses are not mutually exclusive, and each would connect PP_1350 to a different layer of adaptation: metal homeostasis, resource allocation, or regulatory insulation.

Beyond establishing strict methylotrophy, we also explored routes to improve growth performance. ALE under glycine-methanol conditions yielded several evolved clones, among which ^Gly^STC1-215 displayed the fastest growth. This strain carried a mutation in lysC predicted to reduce allosteric regulation, based on analogous substitutions in *E. coli*\ and *Corynebacterium glutamicum* (Liu et al., 2024; Petkevičius et al., 2025), which may increase flux toward glycine regeneration. The consistent co-occurrence of metF mutations suggests coordinated adjustment of a branch point shared with methionine and lysine biosynthesis, potentially rebalancing competing demands on the threonine-glycine node.

Strain ^Gly^STC1-215 also carried mutations in *roxS*. RoxS is an integral-membrane histidine kinase that acts with the response regulator RoxR. In *P. putida* KT2440, RoxSR deletion alters expression of genes involved in amino acid and sugar metabolism, sulfur starvation, respiratory-chain function, and redox homeostasis (Fernández-Piñar et al., 2008). Of direct relevance here, *roxSR* loss is known to upregulate *ltaE, glyA-I*, and *fdhA* (Fernández-Piñar et al., 2008). In *P. putida*, RoxSR has also been linked to cadmium tolerance (Royet et al., 2025) and seed adhesion (Fernández-Piñar et al., 2008), whereas in P. aeruginosa, this regulatory system affects epithelial association and cyanide-insensitive oxidase gene expression (Comolli and Donohue, 2002; Hurley et al., 2010). The *roxS* mutations recovered during ALE may therefore reflect selection for altered redox and energy regulation, but we favor the interpretation that they increase expression of STC-related enzymes and formaldehyde oxidation capacity, thereby reducing toxicity and supporting C_1_ assimilation. This interpretation also fits the recurrent selection of mutations that moderate formaldehyde formation while maintaining downstream detoxification and assimilation capacity.

Overall, we evolved and characterized *P. putida* strains capable of STC-dependent growth under fully methylotrophic conditions, including assimilation of atmospheric CO_2._ ALE overcame several constraints in methanol oxidation, glycine regeneration, transhydrogenase activity, STC enzyme expression, and regulatory balance. In this context, the mutations described here should be viewed as starting points for further design-build-test-learn cycles, where individual alleles can be reconstructed, combined, and tested under increasingly stringent C_1_-dependent regimes. More broadly, this work adds to a growing body of studies showing that synthetic C_1_ assimilation has progressed from pathway design toward functional implementation in heterotrophic hosts. Comparative analyses across organisms will remain essential, because host-specific metabolic architectures strongly shape pathway behavior and evolutionary outcomes (Favoino et al., 2025). In this study, introduction of methylobacterial formate assimilation genes, targeted deletions for growth coupling, and only seven ALE-derived mutations were sufficient to enable strict methanol-dependent growth through the STC in *P. putida*. This compact evolutionary solution contrasts with studies in which hundreds of mutations accumulated during ALE, often through hypermutator phenotypes. At the same time, our findings align with iterative ALE to establish a Calvin-Benson-Bassham cycle in *E. coli*, where as few as three mutations supported effective autotrophic growth (Ben-Nissan et al., 2024). Overall, the findings reported in this article position *P. putida* as a genetically tractable chassis for synthetic methylotrophy and define practical engineering targets for increasing STC flux, specific growth rates and cell densities, and process relevance.

Finally, and from an industrial microbiology perspective, life cycle assessment will be important to determine whether full methylotrophy is required for fueling sustainable industrial processes. Our results demonstrate that co-feeding small amounts of glycine or acetate, which can be potentially derived from plant or microbial biomass or through acetogenesis, enable robust growth under mixotrophic conditions. Acetate is especially relevant because it can be produced from C_1_ gases, including CO and CO_2_, using renewable energy, green H_2_, or syngas. These scenarios may translate into viable processes that remain more sustainable than sugar-or fossil-based alternatives.

## 4. MATERIALS AND METHODS

### 4.1. Chemicals, bacterial strains, medium composition, and culture conditions

Chemicals were purchased from Sigma-Aldrich Co. (St. Louis, MO, USA) unless indicated otherwise. Oligonucleotides were synthesized by Integrated DNA Technologies Inc. (Coralville, IA, USA), and DNA sequencing was performed at Eurofins Genomics Germany GmbH (Ebersberg, Germany). All primers are listed in Table S7. PCR reactions used Phusion *U* Hot Start^TM^ DNA polymerase from Thermo Fisher Scientific Inc. (Waltham, MA, USA), and colony PCRs used OneTaq^TM^ master mix from New England BioLabs Inc. (Ipswich, MA, USA).

All bacterial strains and plasmids are listed in Tables S5 and S6, respectively. E. coli DH5α λpir (Platt et al., 2000) was used as cloning host, whereas the reduced-genome *P. putida* strain SEM11 (Wirth et al., 2023b) was used for quantitative physiology and engineering unless indicated otherwise. Cultivations were performed in lysogeny broth (LB), minimal salt medium (MSM), or modified M9 (mM9) medium as indicated in the text (Green and Sambrook, 2012; Hartmans et al., 1989; Nikel et al., 2008; Pardo et al., 2022). LB contained 10 g L^−1^ tryptone, 5 g L^−1^ yeast extract, and 10 g L^−1^ NaCl. MSM contained 3.88 g L^−1^ K_2H_PO_4_, 1.63 g L^−1^ NaH_2P_O_4_, 2 g L^−1^ (NH_4)2S_O_4_, and 0.1 g L^−1^ MgCl_2·_6H_2O_, with the initial pH adjusted to 7.0. Whenever needed, minimal media was supplemented with a trace element solution that contained 10 mg L^−1^ disodium ethylenediaminetetraacetic acid (EDTA), 2 mg L^−1^ ZnSO_4·_7H_2_O, 1 mg L^−1^ CaCl_2·_2H_2_O, 5 mg L^−1^ FeSO_4·_7H_2_O, 0.2 mg L^−1^ Na_2_MoO_4·_2H_2_O, 0.2 mg L^−1^ CuSO_4·_5H_2_O, 0.4 mg L^−1^ CoCl_2·_6H_2_O, and 1 mg L^−1^ MnCl_2·_2H_2_O. The composition of mM9 medium is described in the ALE section. In some experiments, TraceCERT^TM^ (Supelco Inc., Bellefonte, PA, USA; cat. # 67349), a standard REE mix containing 16 elements (Sc, Y, La, Ce, Pr, Nd, Sm, Eu, Gd, Tb, Dy, Ho, Er, Tm, Yb, and Lu) at 50 mg L^−1^ in HNO_3_, were added to the medium as explained in the text. When required, kanamycin and gentamicin were supplied at 50 μg mL^−1^ and 10 μg mL^−1^, respectively.

Overnight LB cultures were diluted 1:100 into 5 mL MSM containing 20 mM glucose in 50 mL culture tubes and incubated at 30°C and 250 rpm for ca. 18 h. Cells were then washed with carbon-free minimal medium and used to inoculate 96-well microtiter plates containing the carbon source(s) indicated in the text. Antibiotics were included in all growth assays unless strains lacked plasmids. For microtiter plate cultivations, 150 μL cell suspensions at OD_600=_ 0.05 were incubated in an Epoch2 microtiter plate reader (BioTek Instruments Inc., Winooski, VT, USA) with 50 μL mineral oil to prevent evaporation. Microtiter plate readings were calibrated against a tabletop spectrophotometer. Specific growth rate (μ) and lag-phase extension (λ) were calculated with QurvE (www.qurveanalysis.com) using smooth spline fits of growth data (Wirth et al., 2023a).

### 4.2. Construction of deletion and overexpression plasmids

Suicide and overexpression plasmids listed in Table S6 were constructed by USER cloning (Bitinaite et al., 2007). For deletion plasmids, ca. 500 bp regions upstream and downstream of the target locus were amplified with Phusion *U* Hot Start^TM^ DNA polymerase (Thermo Fisher Scientific Inc.) using uracil-containing primers. The pGNW2 backbone was digested with DpnI, after which 1 μL of DpnI-treated vector was mixed with 100 ng of each PCR fragment and 1 μL USER^TM^ enzyme (New England BioLabs Inc.) in a final volume of 10 μL. Reactions were incubated for 30 min at 37°C, cooled from 28°C to 20°C over 3 min at 1°C per step, and held at 10°C for at least 10 min. Chemically competent *E. coli* DH5α λpir cells were transformed by heat shock with 5 μL USER mix, recovered, and plated on selective LB agar.

### 4.3. Construction of mutant P. putida strains

Suicide pGNW2-derivative plasmids were delivered by triparental conjugation using *E. coli* DH5α λpir carrying the suicide plasmid, the recipient *P. putida* strain, and *E. coli* HB101 carrying plasmid pRK2013 as helper strain (Figurski and Helinski, 1979). The three strains were incubated on LB plates for >5 h at 30°C and then plated on LB agar containing the appropriate antibiotic and Irgasan. Positive co-integrants were transformed with plasmid pQURE6·H (Table S6), a conditionally replicative plasmid encoding I-SceI (Volke et al., 2021b). I-SceI cleaves pGNW2 co-integrants in the chromosome and forces a second homologous recombination event. For this step, 50 ng plasmid DNA was electroporated into 50 μL freshly prepared electrocompetent *P. putida* cells washed three times with 300 mM sucrose. Electroporation was performed in 2 mm cuvettes using a Gene Pulser XCell electroporator (Bio-Rad Laboratories Inc., Hercules, CA, USA) set to 2.5 kV, 25 μF, and 200 Ω. Cells were recovered in 1 mL LB supplemented with 2 mM 3-methylbenzoate (3-mBz) for at least 3 h at 30°C and plated on LB agar containing the appropriate antibiotic(s) and 1 mM 3-mBz to induce plasmid replication and I-SceI expression. Positive clones were identified by colony PCR, verified by DNA sequencing, and cured from the resolving plasmid by serial dilution under non-selective conditions.

### 4.4. Parsimonious flux balance analysis

Parsimonious flux balance analysis was performed with the COBRApy Python package to compare STC and wild-type conditions *in silico* (Ebrahim et al., 2013; Heirendt et al., 2019). Simulations used a curated version of the latest genome-scale metabolic model for *P. putida* KT2440 (Bergen et al., 2025; Nogales et al., 2020; Puiggené et al., 2025b). Reactions corresponding to the C_1_-assimilation modules were introduced into the model. Carbon-equimolar uptake rates were set for the respective primary substrates at 36 mmol C g cell dry weight^−1^ h^−1^. Scripts and additional details on the protocols are available at the https://github.com/puiggene07/PubSuppl in the 2024_Ser_Cycles_P_putida directory.

### 4.5. Whole-genome sequencing

*P. putida* genomic DNA was extracted from LB-grown cells using DNeasy Blood & Tissue Kits (QIAGEN GmbH, Hilden, Germany). Samples were sent to Plasmidsaurus Inc. (Eugene, OR, USA) for Nanopore sequencing. Raw reads were assembled against the *P. putida* reference genome using Geneious 8.1 software (Biomatters Ltd., Auckland, New Zealand). Mutations with frequencies >60% were identified by comparative analysis against the corresponding parental strains.

### 4.6. Labelling experiments

Cultures were grown in biological triplicates until stationary phase in MSM supplemented with 20 mM glucose or glycine, 125-250 mM ^13^C-methanol, and 10 nM LaCl_3._ Biomass was harvested, and samples were prepared for GC-MS analysis of isotopomer distributions in proteinogenic amino acids (Long and Antoniewicz, 2019). Fragments containing the full carbon backbone of the respective amino acid were used for further analysis: Ala 260, Gly 246, Val 288, Ser 390, Thr 404, Phe 336, Asp 418, Glu 432, Lys 431, and His 440. Processed data were corrected for natural isotope abundance in the derivatization agents used for GC-MS analysis (Wahl et al., 2004).

### 4.7. RNA sequencing and analysis

Cultures were grown in biological triplicates until mid-exponential phase. Biomass was harvested, washed with phosphate buffer containing 12 mM phosphate buffer, 2.7 mM KCl, and 137 mM NaCl at pH 7.4, and pellets corresponding to 1 mL at OD_600=_ 2 were flash-frozen in liquid N_2 a_nd stored at – 70°C. RNA extraction, purification, and transcriptomics were performed by GENEWIZ Germany GmbH, an Azenta Life Sciences company (Leipzig, Germany). RNA sequencing was used to analyze transcriptomic responses under the carbon-source regimes indicated in the manuscript. Raw data were processed with DESeq2 v1.48.1 in R v4.5.1 using standard pipelines for normalization, differential expression analysis, and significance testing. Differentially expressed genes were identified using DESeq2 default parameters, with *p*-values adjusted by the Benjamini-Hochberg method to control the false discovery rate. Principal component analysis, MA plots, and volcano plots were generated using a modified version of VisomX (available at https://github.com/NicWir/VisomX). The secondary structure and positional entropy analysis of the tRNA for L-threonine encoded by *PP_t73* was generated and analyzed using RNAfold, a ViennaRNA Package (Gruber et al., 2008).

### 4.8. ALE campaigns towards establishing full methylotrophy

The mM9 medium contained 0.011 g L^−1^ CaCl_2_, 0.24 g L^−1^ MgSO_4_, 1× M9 salts, 1× Wolfe’s vitamins, 1× M9 trace elements, 1× SL7 trace elements, and a commercial (TraceCERT^TM^) REE mix. The 10× M9 salts solution contained 5 g L^−1^ NaCl, 20 g L^−1^ (NH_4)2S_O_4_, 30 g L^−1^ KH_2P_O_4_, and 68 g L^−1^ Na_2H_PO_4._ The 2,000× M9 trace element solution contained 15 g L^−1^ disodium EDTA, 4.5 g L^−1^ ZnSO_4·_7H_2_O, 0.7 g L^−1^ MnCl_2·_4H_2_O, 0.3 g L^−1^ CoCl_2·_6H_2_O, 0.26 g L^−1^ CuSO_4·_5H_2_O, 0.4 g L^−1^ Na_2_MoO_4·_2H_2_O, 4.5 g L^−1^_C_aCl_2·_2H_2_O, 3 g L^−1^ FeSO_4·_7H_2_O, 1 g L^−1^ H_3B_O_3_, and 0.1 g L^−1^ KI. The 1,000× Wolfe’s vitamin solution contained 10 mg L^−1^ pyridoxine hydrochloride, 5 mg L^−1^ each of thiamine hydrochloride, riboflavin, nicotinic acid, calcium D-(+)-pantothenate, *p*-aminobenzoic acid, and thioctic acid, 2 mg L^−1^ each of biotin and folic acid, and 1 mg L^−1^ vitamin B_12._ The 1,000× SL7 solution contained 2.1 g L^−1^ FeSO_4·_7H_2_O, 190 mg L^−1^ CoCl_2·_6H_2_O, 100 mg L^−1^ MnCl_2·_4H_2_O, 70 mg L^−1^ ZnCl_2_, 62 mg L^−1^ H_3B_O_3_, 36 _m_g L^−1^ Na_2M_oO_4·_2H_2_O, 28 mg L^−1^ NiSO_4·_6H_2_O, and 32 mg L^−1^ CuSO_4·_5H_2_O. REE were added by diluting 2,000-fold the TraceCERT^TM^ solution. All media were adjusted to pH = 7.0 and filter sterilized.

Five independent ALE replicates were performed on a customized liquid-handler platform as previously described (LaCroix et al., 2015). We used a protocol developed to adapt microorganisms to non-native substrates with support from a low concentration of growth-enabling supplement (Guzmán et al., 2019). In these experiments, mM9 medium contained 500 mM methanol, 17.62 g L^−1^, as the non-native substrate and 5 mM (0.37 g L^−1^) glycine, as the initial growth-enabling supplement. The protocol started with a weaning regime in which glycine concentration was dynamically adjusted to force increasing reliance on methanol. Lineages that adapted to methanol as sole substrate were propagated without supplement. Lineages that did not were maintained under a supplement regime in which glycine was held constant or reduced stepwise until methanol-only growth was achieved.

Growth was monitored periodically by OD_600 u_sing a Tecan Sunrise microplate absorbance reader (Tecan Group Ltd., Männedorf, Switzerland). Cultures were maintained in stationary phase for ca. 48 h after reaching the target OD_600 o_f 0.2 (Fig. S4). Cultures that failed to grow after 3 days were reinoculated with 1 mL from the previous tube. At each passage, stationary-phase cultures were split into a supplemented tube and a methanol-only tube containing 500 mM methanol. Glycine concentration in the next supplemented tube was decreased or increased depending on whether cultures surpassed OD_600=_ 0.2. Methanol-only tubes were considered to grow once OD_600 r_eached at least 0.15, after which 800 μL was propagated to a fresh tube. Tubes without growth after 5 days were discarded. After at least five consecutive methanol-only propagations, weaning was considered successful and cultures were maintained without glycine. Lineages unable to sustain methanol-only growth were evolved under a fixed-supplement regime. Glycine was initially kept at 2.63 mM (0.197 g L^−1^), a concentration sufficient to support faster growth, and later increased to 3.00 mM (0.225 g L^−1^), to further increase growth rates. Cultures were passaged during mid-exponential phase to select for faster-growing phenotypes.

All ALE replicate lineages were initiated from independent colonies. Cultures were grown in vessels containing 17 mL of mM9 medium at 30°C with vigorous stirring at 1,200 rpm. Samples were periodically recovered from spent medium by centrifugation at 1,500×g for 6 min, resuspended in carbon-free mM9 containing 25% (v/v) glycerol, and stored at –70°C for reference and later sequencing.

### 4.9. Data processing and statistical analysis

Data were analyzed using Prism 9.0.2 (GraphPad Software Inc., San Diego, CA, USA). Values are reported as averages ± standard deviation from at least three independent biological replicates, as specified in the corresponding figure legends. Because most negative controls, including strains transformed with empty vectors, showed no growth, variants that restored methylotrophic growth were considered statistically significant (Ferreira, 2007). When relevant, statistical tests are indicated in the corresponding figure legends.

## AUTHOR′S CONTRIBUTIONS

O.P., E.O., and P.I.N. conceived the project and designed the experimental strategy. S.D. performed and analyzed the isotope-labelling experiments. M.F., R.R., and E.T.M. performed the ALE experiments with supervision by A.M.F. L.F.M.J., C.L., S.H.K., V.K., and E.Ö. characterized evolved clones and supported ALE efforts under the supervision of J.F. O.P. and E.Ö. analyzed the data. O.P. performed the remaining experiments leading to the results described in the article. All authors contributed to scientific discussions and interpretation of the results. O.P. and P.I.N. wrote the manuscript, with contributions from all authors.

## ACKNOWLEDGEMENTS

We thank the analytics team, especially Linda Ahonen and Karsten Steen Jensen, for technical support with the amino acid labelling experiments. Financial support to P.I.N. was provided by the Novo Nordisk Foundation through TARGET (NNF21OC0067996), NNF20CC0035580, and BRIGHT (NNF24SA0100980), and by the European Union’s Horizon 2020 Research and Innovation Programme under grant agreement No. 101082049 (TOLERATE). E.O. received support from the European Union through Marie Skłodowska-Curie grant agreement No. 101065339, and from the Novo Nordisk Foundation through UNMUTE (NNF25OC0100601) and GLYCO2 (NNF25OC0103238).

