## Supplementary Information for "Seven mutations unlock strict synthetic methylotrophy in engineered *Pseudomonas putida*"

by

Òscar Puiggené^a^*, Martina Fricano^a^, Riccardo Rossi^a^, Luc F. M. Jansen^a^, Emre Özdemir^a^, Se Hyeuk Kim^a^, Christina Lenhard^a^, Elsayed T. Mohamed^a^, Stefano Donati^a^, Jochen Förster^a^, Viji Kandasamy^a^, Adam M. Feist^a,b^, Enrico Orsi^a^* and Pablo I. Nikel^a^*

^a^	The Novo Nordisk Foundation Biotechnology Research Institute for the Green Transition (BRIGHT), Technical University of Denmark, Kongens Lyngby, Denmark

^b^	Department of Bioengineering, University of California, San Diego, La Jolla, California, USA

**Running title:**	Strict synthetic methylotrophy in *P*. *putida*

**Keywords:**	*Pseudomonas putida*; Synthetic Biology; Synthetic Metabolism; Adaptive Laboratory Evolution; Metabolic engineering; Methanol

***** Correspondence to:

**Òscar Puiggené**

**Enrico Orsi**

**Pablo I. Nikel**

The Novo Nordisk Foundation Biotechnology Research Institute for the Green

Transition (BRIGHT), Technical University of Denmark

2800 Kongens Lyngby, Denmark

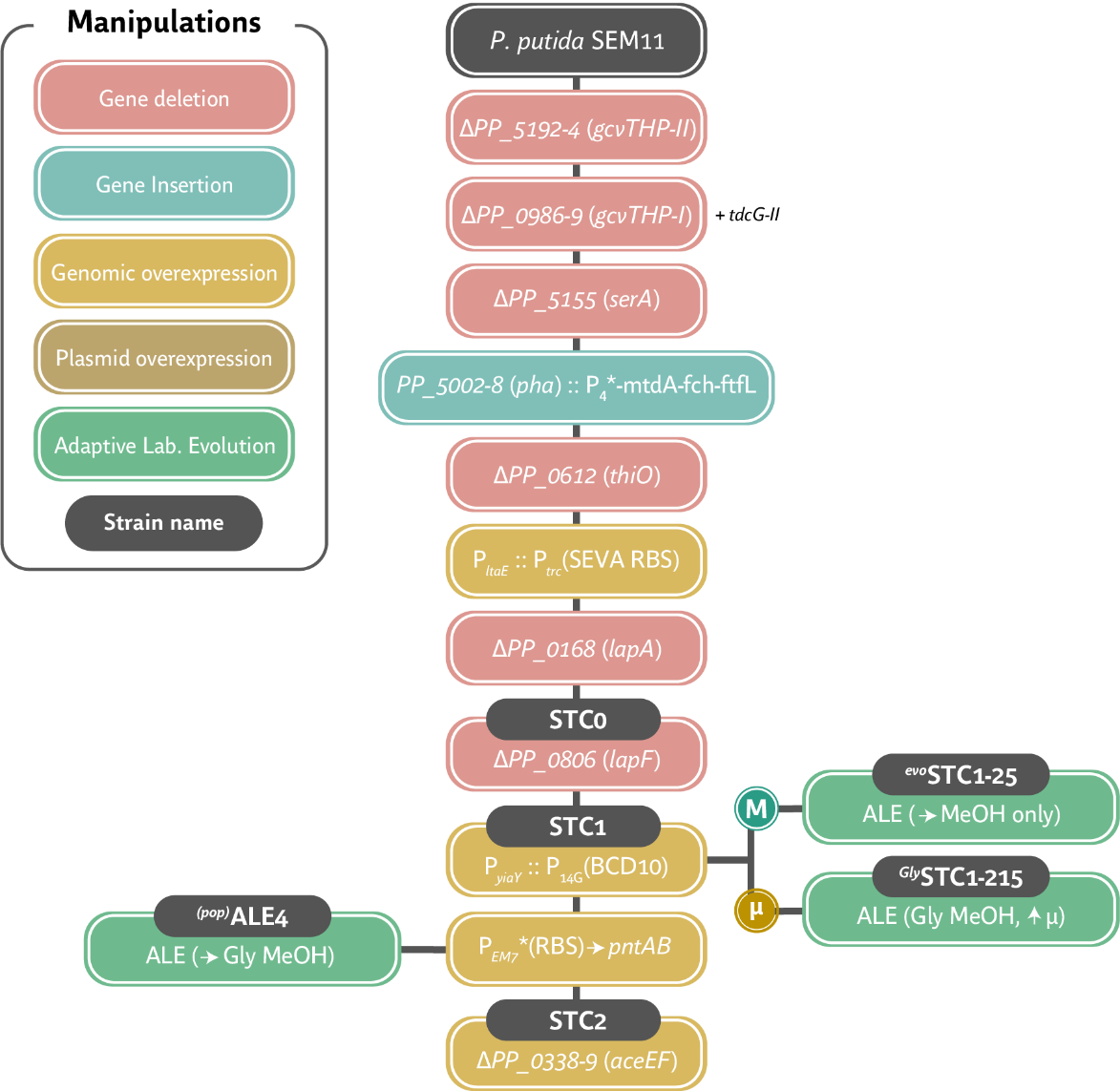

**Figure S1.** Modification chart of the several growth-coupled selection strains constructed and tested in this study.

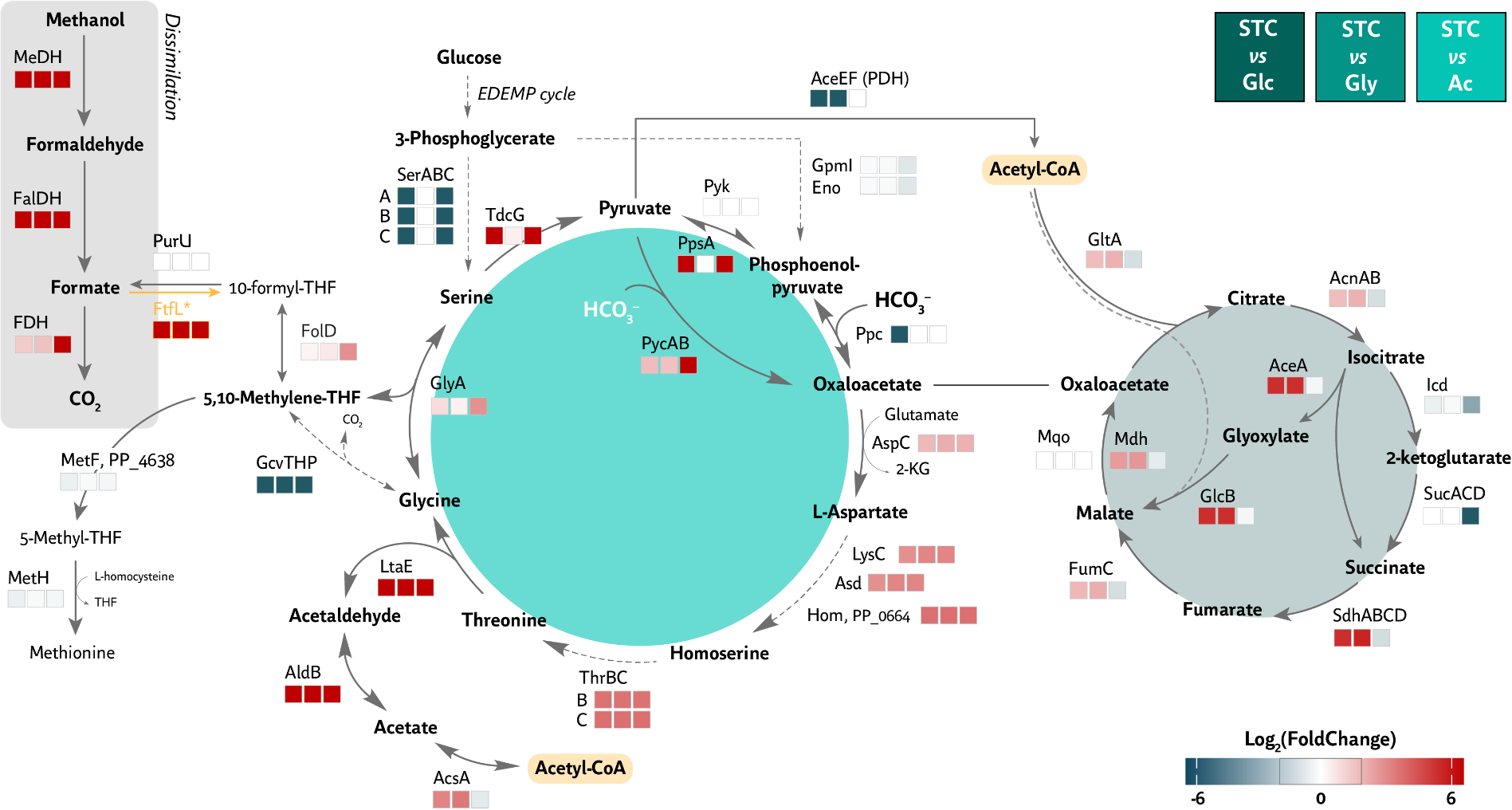

**Figure S2.** pFBA prediction of the fluxes required for optimal growth on the serine-threonine cycle compared to growth on glycose, glycine and acetate. Log_2_ of fold change in flux (mmol/g cell dry weight/h) is given in each case.

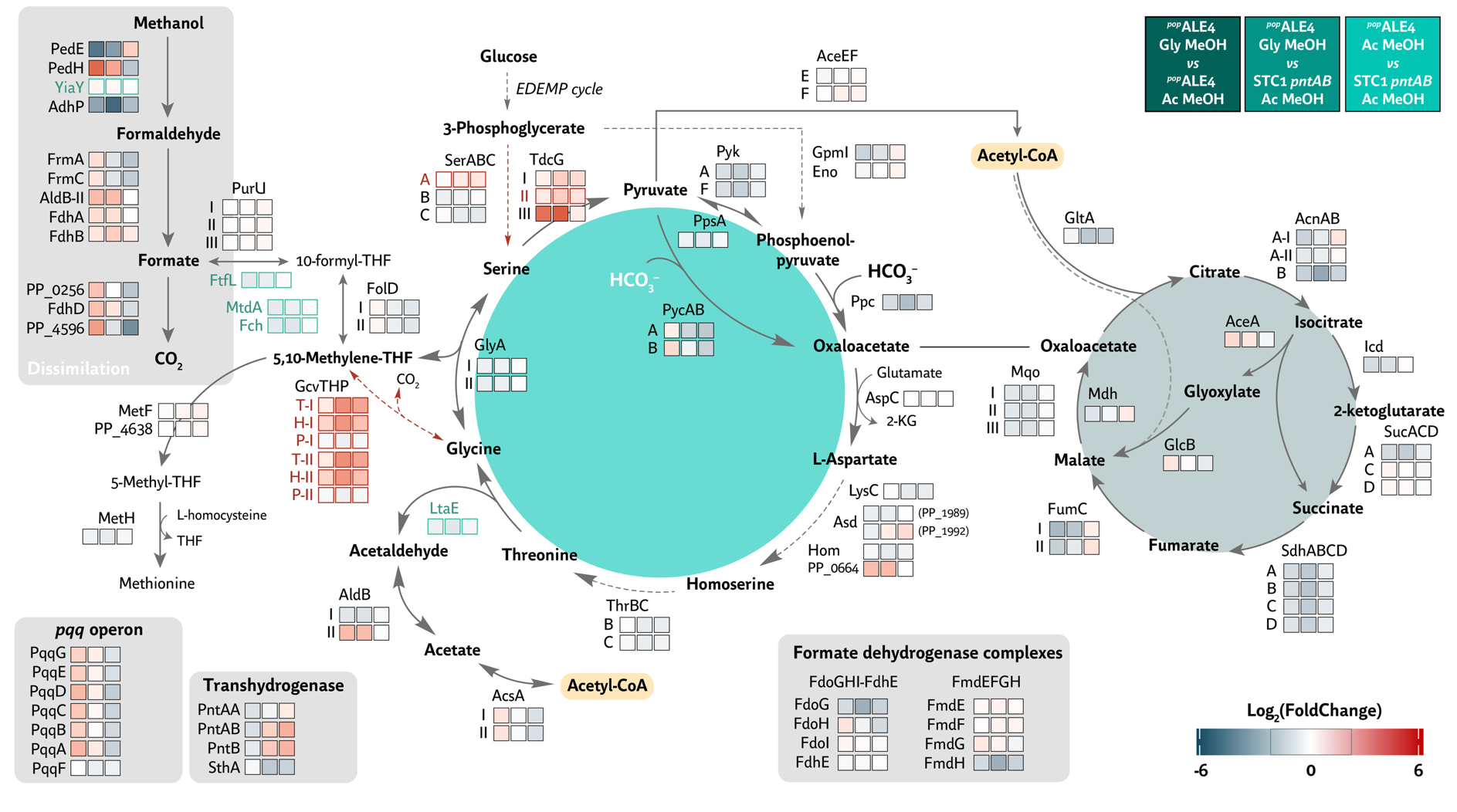

**Figure S3.** Transcriptomics data of population *^pop^*ALE4 in acetate-methanol and glycine-methanol conditions compared to its parental strain STC1 P*_EM7_*® *pntAB* growing on DBM medium with acetate and methanol. Legends are given in the top-right corner of the figure.

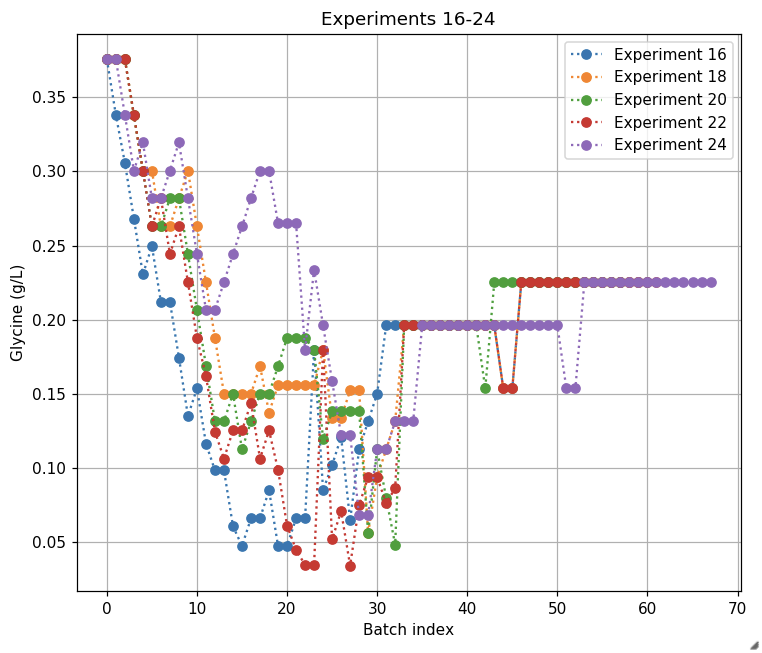

**Figure S4.** ALE strategy to (1) decrease glycine concentration and (2) increase growth rate on this secondary carbon source, with STC1 strain in mM9 and 500 mM MeOH. The concentration of glycine is plotted against the batch number on the right y-axis (g/L) and is represented by different colours depending on the experiment. This concentration demonstrates the progressive step changes in carbon source concentration over the course of the experiment.

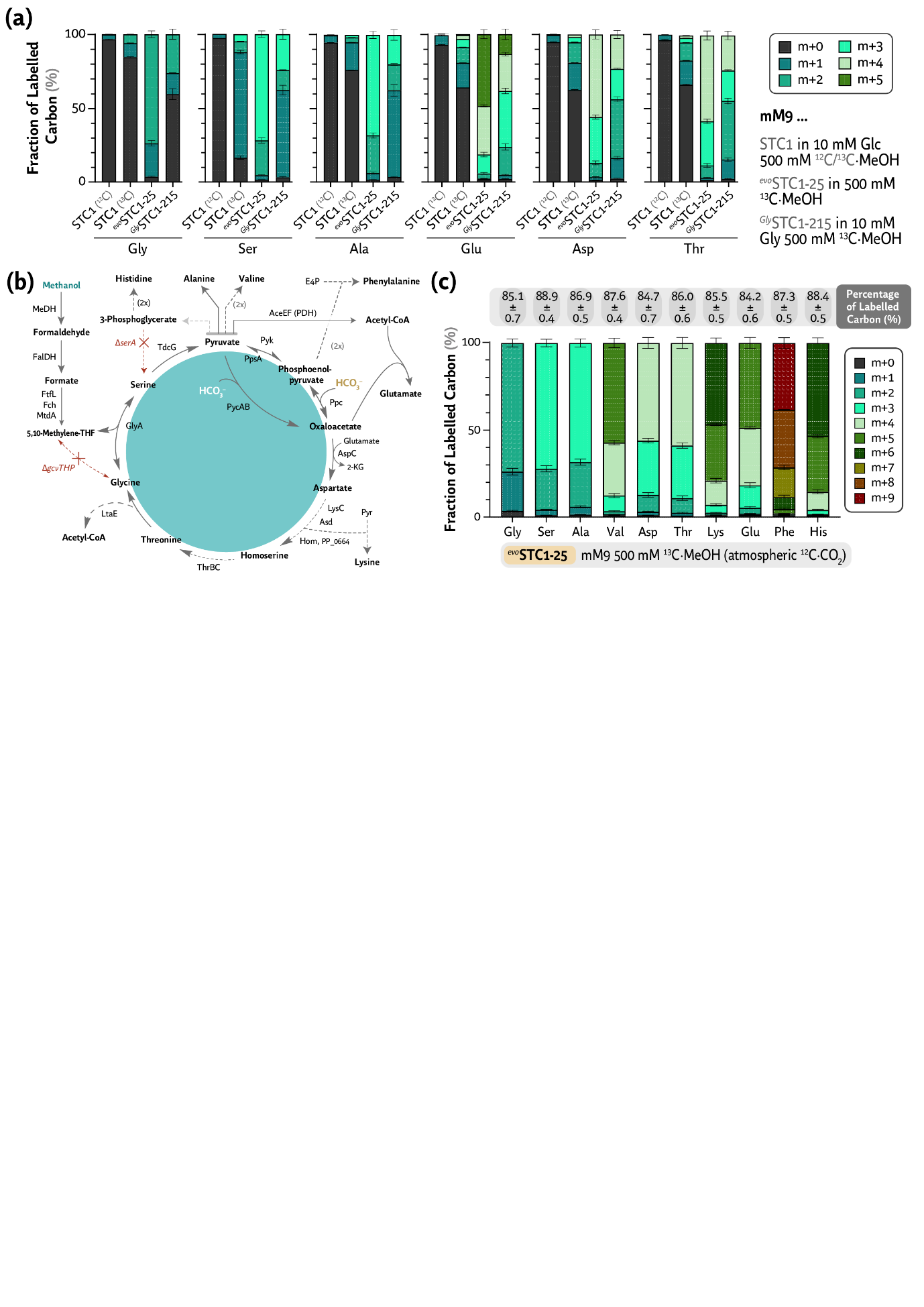

**Figure S5.** **Isotope labelling of *^evo^*STC1-25 and *^Gly^*STC1-215 compared to their parental control STC1**. **(a)** Labelling pattern of amino acids glycine, serine, alanine, glutamate, aspartate and threonine of strains STC1, *^evo^*STC1-25 and *^Gly^*STC1-215 grown in mM9 medium and glucose-methanol, methanol-only or glycine-methanol respectively, as indicated in the legend of the caption. STC1 grown in unlabelled ^12^C-methanol is used as a control. **(b)** Metabolic map of the STC connected to the central carbon metabolism of *P. putida* indicating amino acid biosynthesis pathways. For more information, see **Fig 1a**. **(c)** Labelling pattern of amino acids noted of *^evo^*STC1-25 grown in mM9 medium supplemented with 500 mM ^13^C-methanol. Atmospheric ^12^CO_2_ accounts for the remaining unlabelled carbon (~10-15%) as the STC can assimilate bicarbonate *via* PEP or pyruvate carboxylases. All experiments are supplemented with 10 nM LaCl_3_ to promote PedH activity. Measurements refer to biological duplicates and standard deviations are included.

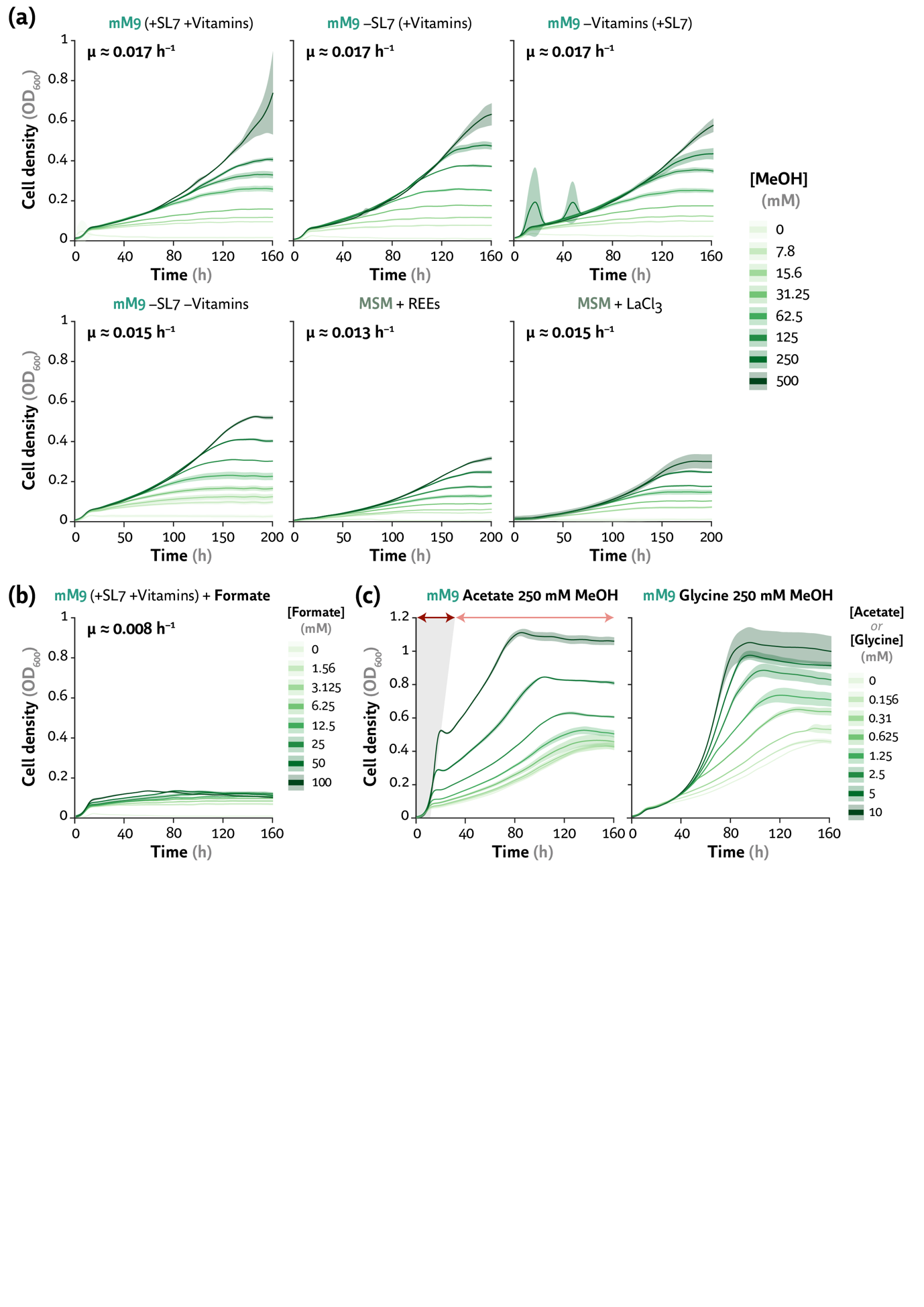

**Figure S6.** *P*. *putida* *^evo^*STC1-25 grown in mM9 or MSM media as a comparison of their growth performance. **(a)** Growth profiles of *^evo^*STC1-25 with or without the presence of SL7 trace elements or Wolfe’s Vitamins (for mM9 medium) or in the presence of REE mix versus 10 nM LaCl_3_ (for MSM) and increasing concentrations of methanol. All mM9 conditions also contained REE mix to promote PQQ-dependent MeDH activity. **(b)** Growth profiles of *^evo^*STC1-25 grown in mM9 medium (with SL7 and Wolfe’s vitamins) and increasing concentrations of sodium formate. **(c)** Growth profiles of *^evo^*STC1-25 grown in mM9, 250 mM methanol and increasing concentrations of acetate (*left*) or glycine (*right*). Final cellular densities and growth rates are reported in **Fig. 2e**.

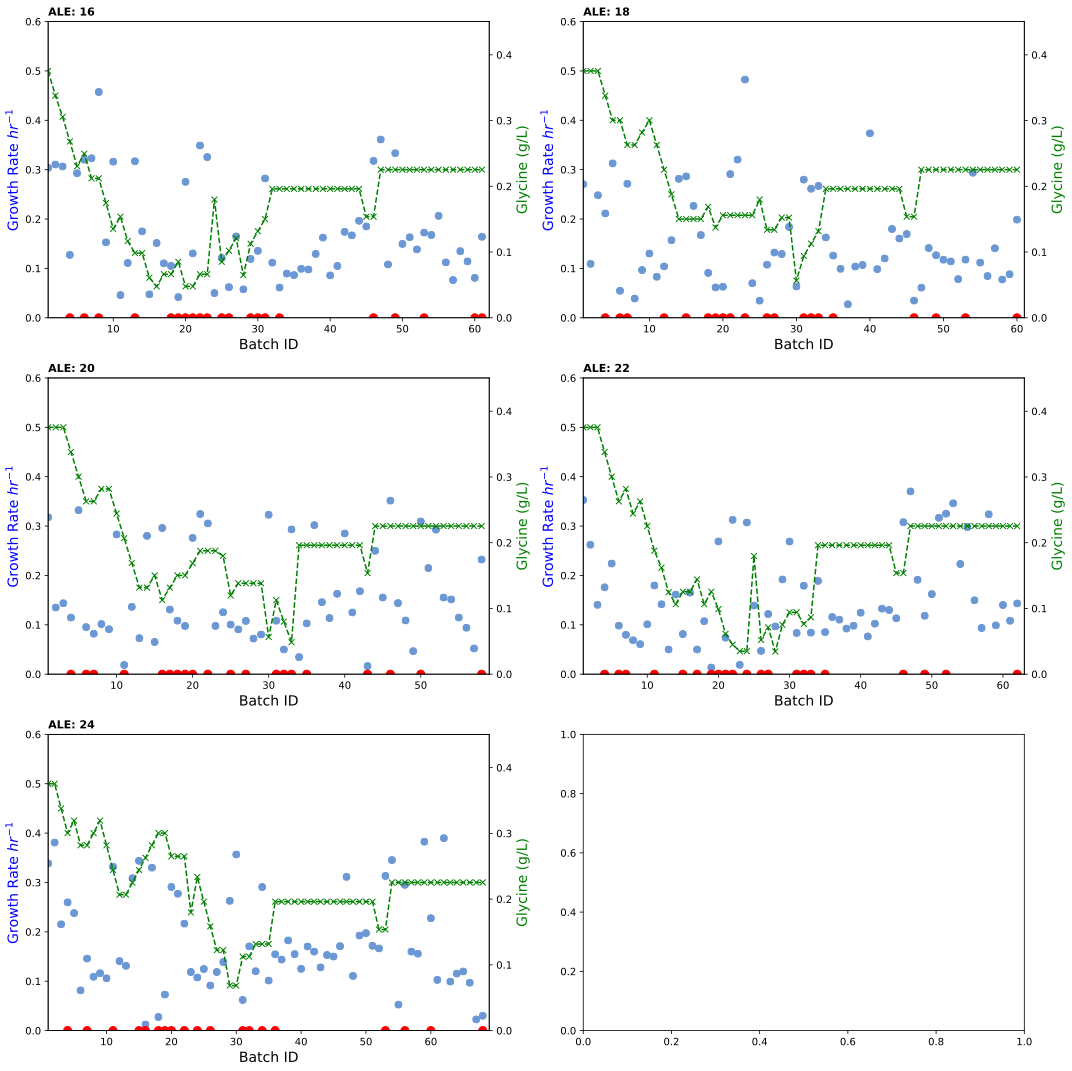

**Figure S7.** ALE experiment fitness trajectories. The population growth rate (h^-1^) for each evolved lineage, presented with their ALE ID, plotted against Batch number. The concentration of glycine is plotted against the batch number in green crosses. Due to the inherent variability of the biological system, including poor growth and repeated culture crashes under the selected conditions, growth rate measurements were inconsistent. The red circle along the *x*-axis represents time-points where cryogenic stock samples of the different linages were saved.

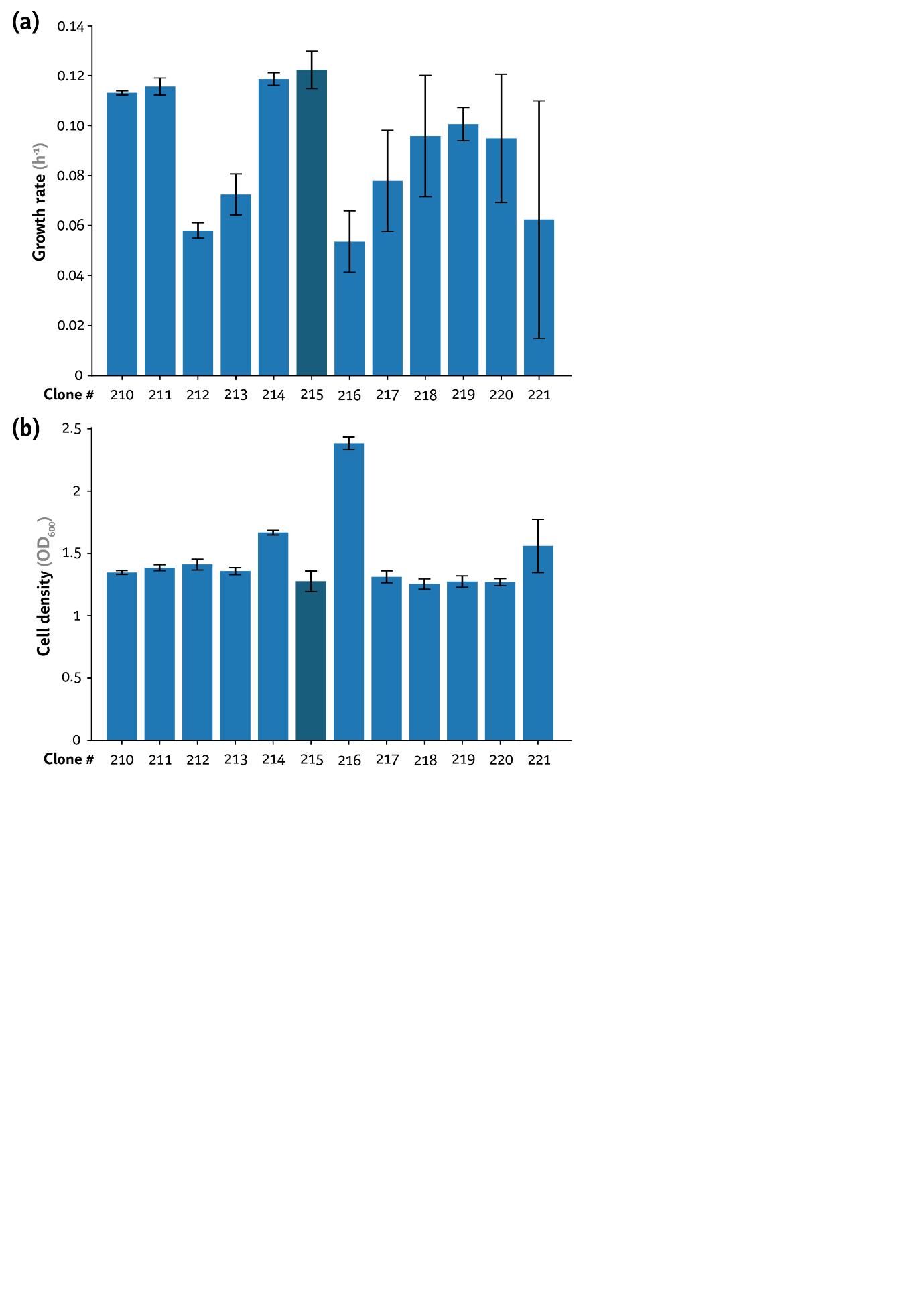

**Figure S8.** Growth parameters of independent clones isolated from the glycine-ALE, towards higher growth rates. Clones were grown in mM9 with 500 mM methanol and 2.7 mM glycine (0.2 g L^-1^) in a growth profiler and growth rate **(a)** and maximum OD_600_ **(b)** are hereby reported. Average values for growth rates (in h^-1^) and bacterial growth (estimated as the optical density measured at 600 nm, OD_600_) ± standard deviation of three biological replicates are represented.

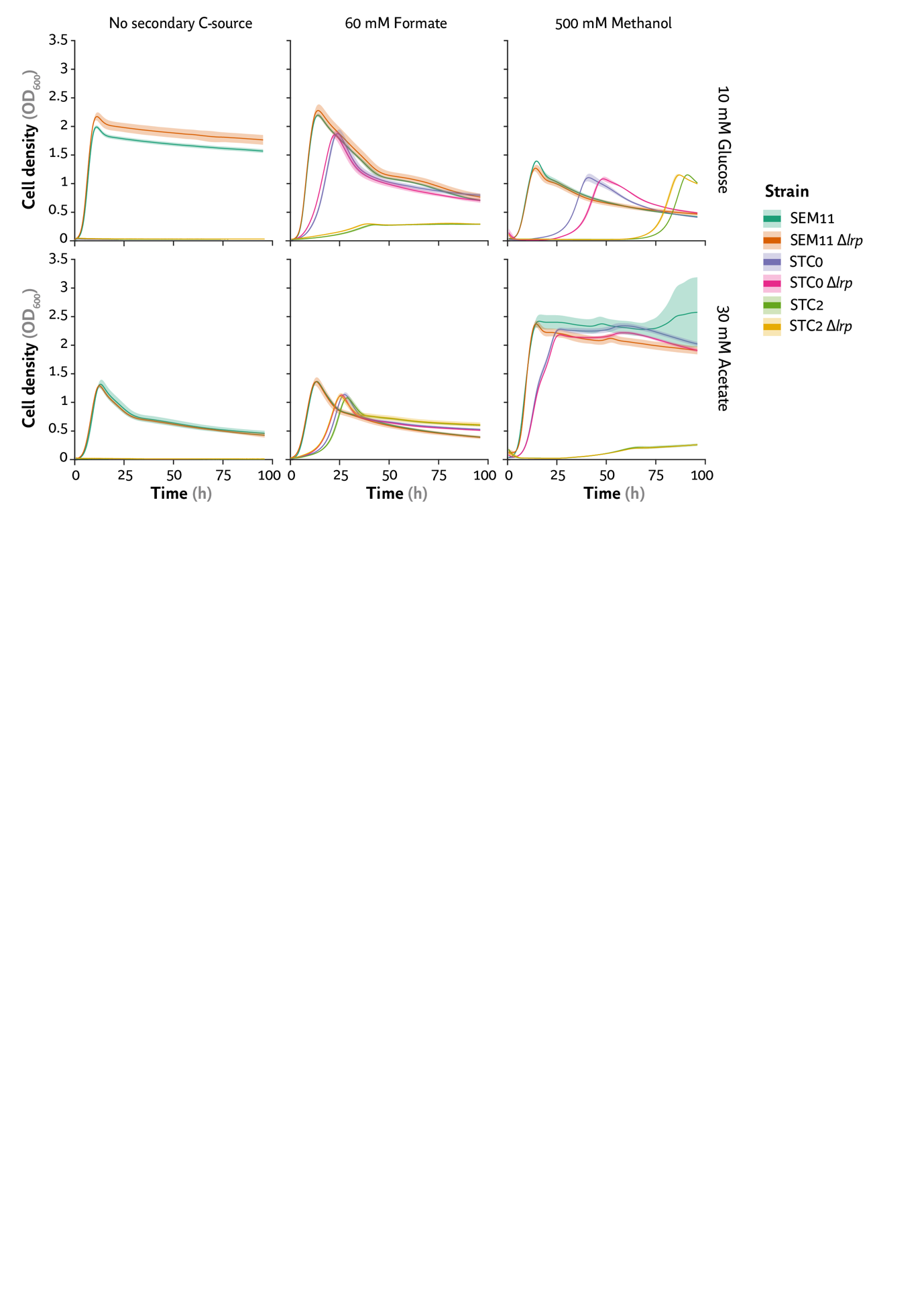

**Figure S9.** Growth profiles comparing strains SEM11, STC0 and STC2 compared to their deletion mutants of gene *lrp*. Cells were grown in MSM supplemented with 10 mM glucose or 30 mM acetate, as well as 60 mM formate or 500 mM methanol (with 10 nM LaCl_3_ for the latter) when indicated. Growth rates have been identified as identical for each WT and ∆*lrp* strains, with only slight differences in lag phase. Average values for bacterial growth (estimated as the optical density measured at 600 nm, OD_600_) ± standard deviation of three biological replicates are represented.

Supplementary Tables

**Table S1.** Enzyme name, function, and reaction for the serine-threonine cycle tested in this study, as well as associated reactions.

| **PP_#** | **Enzyme** | **Function** | **Reaction** |
| --- | --- | --- | --- |
| **Serine-threonine cycle** | | | |
| – | MeDH | Methanol dehydrogenase | NAD + Methanol <=> **NADH** + Formaldehyde |
| PP_1616-7 PP_0328 | FalDH | Formaldehyde dehydrogenase | NAD + Formaldehyde + H_2_O <=> **NADH** + Formate |
| – | FtfL | Formate tetrahydrofolate ligase | **ATP** + Formate + Tetrahydrofolate <=> ADP + Orthophosphate + 10-Formyltetrahydrofolate |
| – | Fch | Methenyltetrahydrofolate cyclohydrolase | 10-Formyltetrahydrofolate <=> 5,10-Methenyltetrahydrofolate + H_2_O |
| – | MtdA | Methylene-tetrahydrofolate synthase | **NADPH** + 5,10-Methenyltetrahydrofolate <=> NADP + 5,10-Methylenetetrahydrofolate |
| PP_0322 PP_0671 | GlyA | Serine hydroxymethyltransferase (SHMT) | Glycine + 5,10-Methylenetetrahydrofolate + H_2_O <=> l-Serine + Tetrahydrofolate |
| PP_0297 PP_0987 PP_3144 | TdcG | Serine dehydratase | l-Serine <=> NH_3_ + Pyruvate |
| PP_5346-7 | PycAB | Pyruvate carboxylase | **ATP** + Pyruvate + CO_2_ <=> ADP + Orthophosphate + Oxalacetate |
| PP_1362 | PykA | Phosphoenolpyruvate synthetase | **ATP** + Pyruvate + H_2_O <=> Orthophosphate + **AMP** + Phosphoenolpyruvate |
| PP_1505 | Ppc | Phosphoenolpyruvate carboxylase | CO_2_ + Phosphoenolpyruvate + H_2_O <=> Orthophosphate + Oxaloacetate |
| PP_3786 | AspC | Aspartate aminotransferase | Oxalacetate + Glutamate <=> l-Aspartate + 2-Ketoglutarate (2-KG) |
| PP_4473 | LysC | Aspartate kinase | **ATP** + l-Aspartate <=> ADP + 4-phospho-l-aspartate |
| PP_1989 PP_1992 | Asd | Aspartate-semialdehyde dehydrogenase | 4-phospho-l-aspartate + **NADPH** <=> l-Aspartate 4-semialdehyde + phosphate + NADP |
| PP_0664 | Hom | Homoserine dehydrogenase | l-aspartate 4-semialdehyde + **NAD(P)H** <=> l-homoserine + NAD(P) |
| PP_0121 | ThrB | Homoserine kinase | Homoserine + **ATP** <=> O-phospho-homoserine + ADP |
| PP_1471 | ThrC | Threonine synthase | O-phospho-homoserine + H_2_O <=> l-Threonine + PO_4_^-^ |
| PP_0321 | LtaE | Low specificity l-threonine aldolase | l-Threonine <=> l-Glycine + Acetaldehyde |
| PP_0545 PP_2680 | AldB | Aldehyde dehydrogenase | Acetaldehyde + **ATP** <=> Acetate + **AMP** |
| PP_4487 PP_4702 | AcsA | Acetyl-coenzyme A synthetase | Acetate + CoA + NAD <=> Acetyl-CoA + **NADH** |
| **Associated reactions to the cycle** | | | |
| PP_0256-7 PP_4596 | Fdh | Formate dehydrogenase | Formate + NAD(P) <=> CO_2_ + **NAD(P)H** |
| PP_0327 PP_1367 PP_1943 | PurU | Formyltetrahydrofolate deformylase | 6R)-10-formyltetrahydrofolate + H2O <=> (6S)-5,6,7,8-tetrahydrofolate + formate |
| PP_1945 PP_2265 | FolD | Bifunctional protein (oxidase and hydrolase of THF moieties) | (6R)-5,10-methylene-5,6,7,8-tetrahydrofolate + **NADP** <=> (6R)-5,10-methenyltetrahydrofolate + **NADPH**  (6R)-5,10-methenyltetrahydrofolate + H_2_O <=> (6R)-10-formyltetrahydrofolate |
| PP_4977  PP_4638 | MetF | Methylenetetrahydrofolate reductase | (6S)-5-methyl-5,6,7,8-tetrahydrofolate + NAD <=> (6R)-5,10-methylene-5,6,7,8-tetrahydrofolate + **NADH** |
| PP_2375 | MetH | Methionine synthase | (6S)-5-methyl-5,6,7,8-tetrahydrofolate + L-homocysteine = (6S)-5,6,7,8-tetrahydrofolate + L-methionine |
| PP_0338 PP_0339 | AceEF | Pyruvate dehydrogenase | Pyruvate + CoA + NAD <=> Acetyl-CoA + CO_2_ + **NADH** |
| **Tricarboxylic acid (TCA) Cycle and Glyoxylate Shunt** | | | |
| PP_4194 | GltA | Citrate synthase | oxaloacetate + acetyl-CoA + H_2_O <=> citrate + CoA |
| PP_2112 PP_2336 PP_2339 | AcnAB | Aconitate hydratase | citrate <=> isocitrate |
| PP_4011 PP_4012 | Icd | Isocitrate dehydrogenase | isocitrate + NADP = 2-oxoglutarate + CO_2_ + **NADPH** |
| PP_4185- PP_4189 | SucABCD | 2-oxoglutarate dehydrogenase | 2-ketoglutarate + NAD + GDP <=> succinate + **NADH** + **GTP** + CO_2_ |
| PP_4190- PP_4193 | SdhABCD | Succinate dehydrogenase | Succinate + Q <=> Fumarate + **QH_2_** |
| PP_0944 PP_1755 | FumC | Fumarate hydratase | Fumarate + H_2_O <=> (S)-malate |
| PP_0751 PP_1251 PP_2925 | Mqo | Malate:quinone oxidoreductase | (S)-malate + Q <=> **QH_2_** + oxaloacetate |
| PP_0654 | Mdh | Malate dehydrogenase | (S)-malate + NAD <=> oxaloacetate + **NADH** |
| PP_4116 | AceA | Isocitrate lyase | D-threo-isocitrate <=> glyoxylate + succinate |
| PP_0356 | GlcB | Malate synthase | glyoxylate + acetyl-CoA + H2O <=> (S)-malate + CoA |
| **Deleted reactions for growth-coupling** | | | |
| PP_5155 | SerA | D-3-phosphoglycerate dehydrogenase | 3-phosphoglycerate + NAD <=> 3-phosphooxypyruvate + **NADH** |
| PP_0986-9 PP_5192-4 | GcvTHP | Glycine cleavage system | 5,10-Methylenetetrahydrofolate + CO_2_ + NADH <=> Glycine + NAD + tetrahydrofolate (THF) |

**Table S2.** Mutation landscape of *P*. *putida* STC1 over the evolution towards low concentrations of glycine in MSM glycine, methanol and lanthanum chloride.

| **Gene** | **Mutations per gene per population** | | | | | | **Function** |
| --- | --- | --- | --- | --- | --- | --- | --- |
|  | **1** | **2** | **3** | **4** | **5** | **6** |  |
| P_14G_(BCD10)  $\to$*yiaY -PP_2683* | – | – | – | – | (BCD10) 46 ∆T | (BCD10) 26 ∆16 nt | RBS of FtfL-Fch-MtdA (i.e. C_1_ module) |
| *yiaY* (*PP_2682*) | – | – | – | C404A (S: P135Q) | – | – | Iron-containing alcohol dehydrogenase, regula-tor of the *ped* cluster |
| *PP_2683* | 814 (T) transposon insertion | – | G446A (S: G149E) | – | – | – | Hybrid histidine kinase, *ped* cluster |
| *pedS1*  *(PP_2664)* | G943T (S: D315Y) | – | – | – | – | – | Hybrid histidine kinase, *ped* cluster |
| *lrp* (*PP_5271*) | – | – | – | G43A (S: D15N) | – | T296G (S: L99R) | Leucine-responsive regulatory protein |
| *PP_5350* | – | 528 ∆G (F) | – | – | – | – | Transcriptional regulator, RpiR family (Glyoxylate shunt repressor) |
| *erdR (PP_1635)* | – | 341 (T) transposon insertion | 341 (T) transposon insertion | – | – | – | DNA-binding response regulator, putatively related to *ped* cluster |
| *mxtR (PP_1695)* | – | – | – | C626T (S: G209D | – | – | Histidine kinase, putatively related to *ped* cluster |
| *cbrB (PP_4696)* | G797A (S: R266H) | C695T (S: A232V) | – | – | – | – | Nitrogen regulation protein NR(I); CbrB; Part of a Two-component system with *PP_4695* |
| *roxS (PP_0887)* | 791+G (F) | – | – | – | – | A350C (S: D117A) | Histidine kinase |
| *roxR* (*PP_0888*) | – | T80G (S: V27G) | – | – | f– | – | Regulatory protein |
| *rpoN* (*PP_0952*) | – | – | – | – | – | G1066A (S: R356C) | RNA polymerase sigma-54 factor |
| *topA* (*PP_2139*) | T1867C (S: C623R) | – | C523T (S: R175C) | – | – | – | DNA topoisomerase |
| *glnD* (*PP_1589*) | – | – | 742 ∆72nt | – | – | – | Bifunctional uridylyl-transferase/uridylyl-removing enzyme |
| *PP_2115* | – | – | – | A130G (S: K44E) / 136 ∆6nt | – | 52 ∆6 nt | Uncharacterised protein |
| *dadA-II* (*PP_5270*) | – | – | – | – | 764 ∆33 nt | – | D-amino acid dehydrogenase |

***Abbreviations****: S: Substitution, T: Truncation; F: Frameshift, ∆: deletion, followed by the size (if not single nucleotide, then the base affected); nt: nucleotide. The amino acid change is indicated between brackets.* Positions at which truncations or deletions occur are given from start codon. Functional interrelatedness of genes is depicted as a colour coding.

**Table S3. Mutation analysis of STC1-25 methylotrophic clones.** 26 clones were whole-genome sequenced and mutations occurring are displayed in here. For clarity, 20 of these clones possessed the same mutations and have been included as representation of *^evo^*STC1-25. 6 differing clones are also included, alongside the genes, their functionality and the mutations reported.

| **Gene** | **Function** | **Mutation** | **Count** | ***^evo^*STC1-25 (20 clones)** | **STC1-25 cl 3** | **STC1-25 cl 4** | **STC1-25 cl 6** | **STC1-25 cl 7** | **STC1-25 cl 27** | **STC1-25 cl 46** |
| --- | --- | --- | --- | --- | --- | --- | --- | --- | --- | --- |
| ***lrp*** | Leucine-responsive regulatory protein | D12Y (GAC>TAC) | 26 | 20 (✓) | ✓ | ✓ | ✓ | ✓ | ✓ | ✓ |
| **P*_pntAB_*** | Transhydrogenase | SNP (-117, C>T) | 26 | 20 (✓) | ✓ | ✓ | ✓ | ✓ | ✓ | ✓ |
| ***ltaE*** | L-threonine aldolase | A73T (GCA>ACA) | 26 | 20 (✓) | ✓ | ✓ | ✓ | ✓ | ✓ | ✓ |
| ***PP_1350*** | Histidine kinase | T261P (ACC>CCC) | 26 | 20 (✓) | ✓ | ✓ | ✓ | ✓ | ✓ | ✓ |
| ***PP_2683*** | Hybrid histidine kinase | Insertion (17, +A) | 26 | 20 (✓) | ✓ | ✓ | ✓ | ✓ | ✓ | ✓ |
| ***PP_t73 (tRNA^Thr^)*** | tRNA for L-threonine | SNP (12, A>G) | 25 | 20 (✓) | ✓ | ✓ | ✓ |  | ✓ | ✓ |
| ***ycgB*** | Regulatory protein with yeaGH | Insertion (394 nt) ((CGGAACAGGTA)1>2) | 25 | 20 (✓) | ✓ | ✓ | ✓ |  | ✓ | ✓ |
| *yeaG* | Serine/threonine-protein kinase | Y463* (TAT>TAA) | 1 |  |  |  |  | ✓ |  |  |
| *PP_1391* | Transcriptional regulator, LysR family | E182D (GAG>GAT) | 1 |  |  |  | ✓ |  |  |  |
| *PP_1421* | Histidine kinase | Insertion (84, (ATGCTG)1>2) | 1 |  |  |  |  | ✓ |  |  |
| *PP_1795, PP_1797* | Uncharacterized protein / Membrane fusion protein (MFP) family protein | Deletion intergenic (-862/-1472, (G)9>8) | 1 |  | ✓ |  |  |  |  |  |
| *PP_2115* | Uncharacterized protein | Insertion (117, (AAGATC)11>12) | 1 |  |  |  |  |  |  | ✓ |
| *PP_2673, qedH-I* | Pentapeptide repeat family protein / PQQ-dependent alcohol dehydrogenase | Deletion intergenic (-38/+29, ∆1 bp) | 1 |  |  |  |  | ✓ |  |  |
| *PP_5606* | Cytochrome D1 | Y314C (TAC>TGC) | 1 |  |  |  |  | ✓ |  |  |
| *PP_3641, PP_3643* | Permease family protein, Oxidoreductase | SNP intergenic (-187/-130, C>A) | 1 |  |  | ✓ |  |  |  |  |
| *PP_5191* | Uncharacterized protein | M75I (ATG>ATA) | 1 |  |  |  |  |  | ✓ |  |

**Table S4. Mutation analysis of STC1 clones evolved towards higher growth rates in glycine-methanol conditions.** 12 clones were whole-genome sequenced and mutations occurring are displayed in here, including the further characterized *^Gly^*STC1-215 (highlighted in yellow).

| **Gene** | **Details** | **Count** | **210** | **211** | **212** | **213** | **214** | **215** | **216** | **217** | **218** | **219** | **220** | **221** |
| --- | --- | --- | --- | --- | --- | --- | --- | --- | --- | --- | --- | --- | --- | --- |
| **P*_pntAB_*** | SNP (-117, C>T) | 4 |  |  | ✓ |  | ✓ | ✓ | ✓ |  |  |  |  |  |
|  | Deletion (-69, ∆5 bp) | 5 |  |  |  |  |  |  |  | ✓ | ✓ | ✓ | ✓ | ✓ |
| ***lrp*** | D12Y (GAC>TAC) | 7 | ✓ | ✓ | ✓ | ✓ | ✓ | ✓ | ✓ |  |  |  |  |  |
|  | R141P (CGC>CCC) | 6 |  | ✓ |  |  |  |  |  | ✓ | ✓ | ✓ | ✓ | ✓ |
| *PP_5350* | S183P (TCC>CCC) | 1 | ✓ |  |  |  |  |  |  |  |  |  |  |  |
|  | Frameshift (486, (G)5>6) | 2 |  | ✓ |  | ✓ |  |  |  |  |  |  |  |  |
| *ltaE* | Y134N (TAC>AAC) | 2 |  |  |  |  | ✓ |  | ✓ |  |  |  |  |  |
| *roxS (PP_0887)* | H121R (CAC>CGC) | 4 | ✓ | ✓ |  | ✓ |  | ✓ |  |  |  |  |  |  |
| *lysC* | G275S (GGC>AGC) | 3 | ✓ |  |  | ✓ |  | ✓ |  |  |  |  |  |  |
| *metF* | L221R (CTG>CGG) | 1 |  |  |  |  | ✓ |  |  |  |  |  |  |  |
|  | G254D (GGC>GAC) | 3 | ✓ |  |  | ✓ |  | ✓ |  |  |  |  |  |  |
| *metW* | A167V (GCC>GTC) | 1 |  |  | ✓ |  |  |  |  |  |  |  |  |  |
| *PP_1350* | Deletion (∆220 bp) | 4 |  |  |  |  |  |  |  | ✓ | ✓ | ✓ |  | ✓ |
|  | Q624L (CAG>CTG) | 2 |  |  |  |  |  |  |  |  |  | ✓ |  | ✓ |
|  | I622T (ATC>ACC) | 5 |  |  |  |  |  |  |  | ✓ | ✓ | ✓ | ✓ | ✓ |
| *agmR* | A189V (GCC>GTC) | 1 |  |  | ✓ |  |  |  |  |  |  |  |  |  |
| P*_yiaY_* | Deletion (-43, ∆1bp) | 1 |  | ✓ |  |  |  |  |  |  |  |  |  |  |
|  | Deletion (-40, ∆1bp) | 2 |  |  |  |  | ✓ |  | ✓ |  |  |  |  |  |
|  | Insertion (-29, (CGTACTGAAAC)1>2) | 5 |  |  |  |  |  |  |  | ✓ | ✓ | ✓ | ✓ | ✓ |
|  | Deletion (-6, ∆9 bp) | 3 | ✓ |  |  | ✓ |  | ✓ |  |  |  |  |  |  |
| *yiaY* | R23W (CGG>TGG) | 6 |  | ✓ |  |  |  |  |  | ✓ | ✓ | ✓ | ✓ | ✓ |
| *PP_2683* | Insertion (17, +A) | 1 |  |  | ✓ |  |  |  |  |  |  |  |  |  |
| *trkA* | G238D (GGC>GAC) | 2 |  | ✓ | ✓ |  |  |  |  |  |  |  |  |  |
| P*_yeaG_* | SNP (-48, G>T) | 1 |  |  |  |  |  |  | ✓ |  |  |  |  |  |
| *acoB* | A171S (GCG>TCG) | 1 |  |  |  |  |  |  |  |  | ✓ |  |  |  |
| *gltI* | S64* (TCG>TAG) | 1 |  |  |  |  |  |  |  |  |  |  |  | ✓ |
| *PP_1795, PP_1797* | Insertion intergenic (-862/-1472, (G)9>10) | 1 |  |  |  |  | ✓ |  |  |  |  |  |  |  |
| *PP_2590* | G390D (GGC>GAC) | 1 |  |  | ✓ |  |  |  |  |  |  |  |  |  |
| *PP_3526* | D78N (GAC>AAC) | 1 |  |  |  |  |  |  |  |  |  | ✓ |  |  |
| *PP_4482* | D228G (GAC>GGC) | 1 |  |  |  |  |  |  | ✓ |  |  |  |  |  |
| *PP_4582, PP_4583* | Deletion intergenic (-54/-29, ∆98 bp) | 1 |  | ✓ |  |  |  |  |  |  |  |  |  |  |
| *folM* | Y226H (TAC>CAC) | 1 |  |  |  |  |  |  |  |  | ✓ |  |  |  |
| *topA* | C692F (TGT>TTT) | 1 |  |  |  |  |  |  | ✓ |  |  |  |  |  |

**Table S5.** Bacterial strains used in this study.

| **Strain** | **Relevant characteristics**^a^ | | **Reference or source** |
| --- | --- | --- | --- |
| *Escherichia coli* | | | |
| DH5α λ*pir* | Cloning host; F^–^ λ*^–^ endA1 glnX44(AS) thiE1 recA1 relA1 spoT1 gyrA96*(Nal^R^) *rfbC1 deoR nupG* Φ*80(lacZ*∆*M15)* ∆*(argF-lac)U169 hsdR17(r_K_^–^ m_K_^+^)*, λ*pir* lysogen | | (Platt et al., 2000) |
| HB101 pRK2013 | Conjugative helper strain; *hsdR-M^+^, proA2, leuB6, thi-1, recA*; harboring plasmid pRK2013, Kan^R^ | | (Figurski and Helinski, 1979) |
| *Pseudomonas putida* | | | |
| KT2440 | Derivative of *P. putida* mt-2 strain, with the TOL plasmid cured (wild-type control) | (Bagdasarian et al., 1981) | |
| KT2440 ∆*PP_5350* | Derivative of *P. putida* KT2440, ∆*PP_5350* | (Volke et al., 2021) | |
| SEM11 | Reduced-genome derivative of *P. putida* KT2440 (Belda et al., 2016) and EM42 (Martínez-García et al., 2014) | (Wirth et al., 2023) | |
| SEM11 ∆*lrp* | Derivative of *P. putida* SEM11, *∆lrp* (*PP_5271*) | This work | |
| STC0 | Derivative of *P. putida* SEM11, *∆serA, ∆gcvTHP-I* (*PP_0986-9*) *∆gcvTHP-II* (*PP_5192-4*) *pha* (*PP_5002-8*)::P_4_*→*mtdA-fch-ftfL* (Turlin et al., 2022)), P*_ltaE_*::P*_trc_* (SEVA RBS) ∆*thiO* (*PP_0612*) ∆*lapA* (*PP_0168*) ∆*lapF* (*PP_0806*) | (Puiggené et al., 2025; Turlin et al., 2022) | |
| STC0 ∆*lrp* | Derivative of *P. putida* STC0, *∆lrp* (*PP_5271*) | This work | |
| STC1 | Derivative of *P. putida* STC0, P*_yiaY_*::P_14G_(BCD10) | (Puiggené et al., 2025) | |
| *^evo^*STC1-25 | Evolved clone from STC1, ALEd towards full methylotrophy. Mutations are included in **Table S2**, alongside other clones differing slightly from this | This work | |
| *^Gly^*STC1-215 | Evolved clone from STC1, ALEd for higher growth rates in glycine-methanol. Mutations are included in **Table S3**, alongside other clones differing from this clone | This work | |
| STC1 P_EM7_*®*pntAB* | Derivative of STC1 harbouring a constitutive, engineered P_EM7_ promoter with increased expression of *pntAB* operon | This work | |
| *^(pop)^*ALE4 | Evolved population from STC1 P_EM7_*®*pntAB* . Clones isolated from the population (*^pop^*) as ALE4 cl#. | This work | |
| STC2 | Derivative of STC1 P_EM7_*®*pntAB*, ∆*aceEF* (*PP_0338-39)* | This work | |
| STC2 ∆*lrp* | Derivative of *P. putida* STC0, *∆lrp* (*PP_5271*) | This work | |

**Table S6.** Plasmids used in this study.

| **Plasmid** | **Relevant characteristics^a^** | **Reference or source** |
| --- | --- | --- |
| pGNW2 | Suicide vector used for deletions in Gram-negative bacteria; *oriT*, *traJ*, *lacZ*α, conditional RK6 replication origin, P_EM7_→*msfGFP*; Km^R^ | (Wirth et al., 2020) |
| pGNW2·∆*lrp* | Derivative of vector pGNW2 carrying homology regions to delete *lrp* (*PP_5271*); Km^R^ | This work |
| pGNW2·∆*PP_5350* | Derivative of vector pGNW2 carrying homology regions to delete *PP_5350*; Km^R^ | This work |
| pGNW2· P_EM7_*→*pntAB* | Derivative of vector pGNW2 carrying homology regions to integrate constitutive promoter *P_EM7_**→*pntAB* as to express *pntAB* operon. Mutation of P_EM7_ via evolution was previously reported in (Turlin et al., 2025); Km^R^ | This work |
| pGNW2·∆*aceEF* | Derivative of vector pGNW2 carrying homology regions to delete *aceEF* (*PP_0338-9)*; Km^R^ | (Wirth et al., 2022) |
| pQURE6·H | Helper plasmid for gene deletions; conditionally-replicating vector carrying XylS/*Pm*→*I-SceI* and P_14G_(BCD2)→*mRFP*; Gm^R^ | (Volke et al., 2020) |
| pS621·*eV* | Control vector derivative of pSEVA621 (Silva-Rocha et al., 2013) harboring a *P_trc_* promoter and the canonical SEVA ribosome binding site (RBS) followed by a start and stop codon (ATGTAA); *oriT oriV*(RK2); Gm^R^ | (Puiggené et al., 2025) |
| pS621·*^Ec^ltaE** (C188Y) | Derivative of vector pSEVA621 harboring a *P_trc_* promoter and the canonical SEVA ribosome binding site (RBS)→ *^Ec^ltaE** (C188Y) (Schann et al., 2024); *oriT oriV*(RK2); Gm^R^ | (Puiggené et al., 2025) |
| pS221·P*_pntAB_*:*msfGFP* | Derivative of vector pSEVA221 harboring the native promoter and RBS of *pntAB* (P*_pntAB_*) expressing *msfGFP*; oriT oriV(RK2); Km^R^ | This work |
| pS221·P*_pntAB_**:*msfGFP* | Derivative of vector pSEVA221 harboring the evolved promoter and RBS of *pntAB* (P*_pntAB_*), with -117C>T mutation, expressing *msfGFP*; oriT oriV(RK2); Km^R^ | This work |
| pS221·*^Pp^ltaE* (WT) | Derivative of vector pSEVA221 harboring the P_14G_ and BCD10 constitutively expressing *ltaE* of *P. putida* (PP_0321); oriT oriV(RK2); Km^R^ | (Puiggené et al., 2025) |
| pS221·*^Pp^ltaE* (A73T) | Derivative of vector pS221·*^Pp^ltaE* (WT), harboring A73T *^Pp^lta*E; oriT oriV(RK2); Km^R^ | This work |
| pS221·*^Pp^ltaE* (Y134N) | Derivative of vector pS221·*^Pp^ltaE* (WT), harboring Y134N *^Pp^lta*E; oriT oriV(RK2); Km^R^ | This work |
| pS221·*^Pp^ltaE* (A73T Y134N) | Derivative of vector pS221·*^Pp^ltaE* (WT), harboring both A73T and Y134N *^Pp^lta*E; oriT oriV(RK2); Km^R^ | This work |
| pMC1·P*_pedE_*:*msfGFP* | Derivative of vector pMC1 vector harboring the native promoter and RBS of *pedE* (P*_pedE_*) expressing *msfGFP*; oriT oriV(RSF1010); Sm^R^ | (Puiggené et al., 2025) |

^a^ Antibiotic markers: Gm, gentamicin; Km, kanamycin; Sm, streptomycin.

**Table S7.** Oligonucleotides used in this study.

| **Name** | **DNA sequence (5’→3’)** | **Use** |
| --- | --- | --- |
| pGNW-USER-fw | AGTCGACCUgcaggcatgcaagcttct | Construction of pGNW2 *plasmids* |
| pGNW-USER-rev | AGGATCUagaggatccccgggtaccg |  |
| ΔPP_5350_Up_fw | AGATCCUGGAGCTACCGCTGTACCGTAGCT | Construction of  pGNW2·∆*PP_5350* |
| ΔPP_5350_Up_rev | ATTCAACCUGTACTGGCCTGTTCGCGGGC |  |
| ΔPP_5350_down_fw | AGGTTGAAUTCACACAGGCGGGACTCGGTTATGA |  |
| ΔPP_5350_down_rev | AGGTCGACUCCGGCAATGCGCAGGTGGTT |  |
| pGNW2-PEM7*-pnt_U_fw | ATGCCGAUGATTAATTGTCAACAGACTATTTC | Construction of  pGNW2· PEM7*→pntAB |
| pGNW2-PEM7*-pnt_rv | ATCGGCAUAGTATAATACGACAAAAGCT |  |
| ∆lrp_Up_fw | AGATCCUCAGTCACGGCTGTACTCGG | Construction of  pGNW2·∆*lrp* |
| ∆lrp_Up_rv | AGCCTCACAUAGGGGATGCCCCTCCGTG |  |
| ∆lrp_down_fw | ATGTGAGGCUAGACCAGCACCTGGCGGG |  |
| ∆lrp_down_rv | AGGTCGACUGCTGCGCGCTGTGGATCA |  |
| pS221-PpntAB:msfgfp_fw | ATAGTCAUGCGTAAAGGTGAAGAACT | Construction of  pS221·P*_pntAB_*:*msfGFP* |
| pS221-PpntAB:msfgfp_rv | AGGGTTAAUTAAAGGCATCAAATAAAACG |  |
| PpntAB_pS221-msfgfp_fw | ATTAACCCUGTAGTCAATTATTTTAAACACCTA |  |
| PpntAB_pS221-msfgfp_rv | ATGACTAUTTCTCCTGCGGTGACCTT |  |
| PpntAB_evomut_pS221-msfgfp_fw | AGCATTTUACAGACGAACCCGAAGGCC | Construction of  pS221·P*_pntAB_**:*msfGFP* |
| PpntAB_evomut_pS221-msfgfp_rv | AAAATGCUTCGCCCCGCCCGGGATCC |  |
| PpltaE A73T_Up_rv | AGGGATGUCAGGGCCAGGGAGTTG | Construction of  pS221·*^Ppl^taE* (A73T) |
| PpltaE A73T_Down_fw | ACATCCCUGTGCCAGAGCTATCACAGCGTGAT |  |
| PpltaE Y134N_Up_rv | ATATCCUGGCGTTTGAGCGCCACTT | Construction of  pS221·*^Ppl^taE* (Y134N) |
| PpltaE Y134N_Down_fw | AGGATAUCCACAACCCCAAGCCG |  |
